# Systematic mapping of human tissue microanatomy reveals age-associated remodeling and resilience

**DOI:** 10.1101/2025.10.13.682110

**Authors:** Iva Buljan, Zsuzsanna Bagó-Horváth, André F. Rendeiro

## Abstract

Aging disrupts tissue structure at various scales, from cellular alterations to tissue and organ-level integrity. Microanatomical domains - recurrent cellular arrangements essential to organ-specific function, provide a highly physiologically relevant perspective on tissue homeostasis but are severely understudied in human aging.

To address this gap, we developed H&E-UTAG, an unsupervised algorithm to detect microanatomical domains in whole slide histopathological images, which enables large-scale, label-free analysis of human microanatomy. Applying it to 24,945 whole-slide images from 40 human tissues of 983 individuals aged 20 to 70 years old, we identified 218 recurrent microanatomical domains categorized into 74 types across tissues. Domain types varied widely in tissue specificity, with 16% shared across 3 or more tissues and 69% restricted to a specific tissue. Age emerged as the dominant factor in influencing domain abundance, with 28% of domains changing significantly over the adult lifespan. By integrating tissue-level pathology annotations, we distinguished structural changes associated with healthy aging from those linked to subclinical disease, revealing that these processes often remodel distinct tissue compartments. Finally, mapping higher-order networks of domain-domain interactions uncovered age-associated reorganization of organ architecture, while a core framework of interactions remained resilient with age.

Our novel analytical framework reveals fundamental principles of tissue organization and how they are restructured across the human lifespan, offering new insights into aging biology and tissue architecture in health and in the path to age-associated diseases.

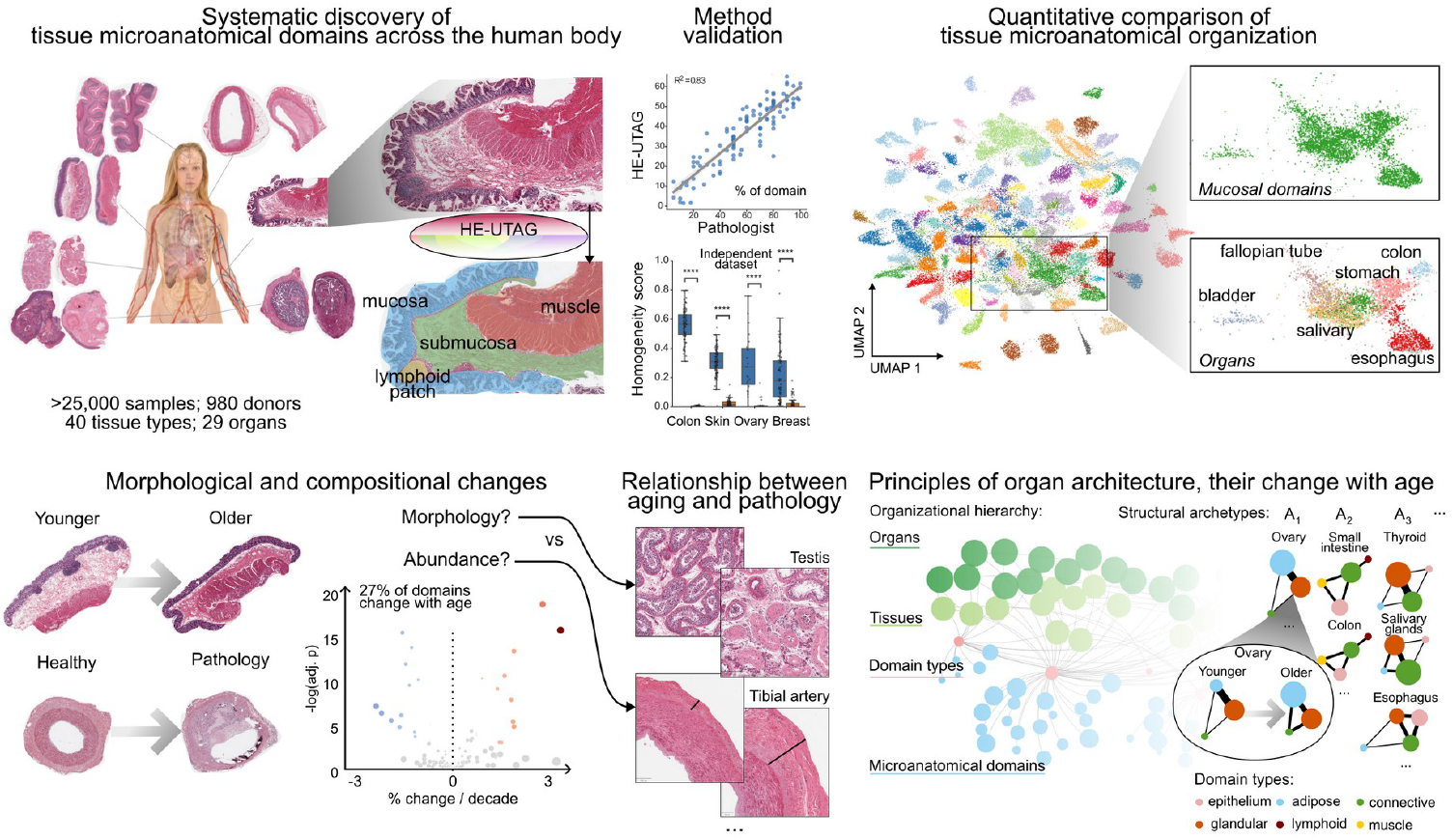

## Introduction

Aging drives structural and functional changes across organs, often manifesting as visible traits such as skin wrinkling, reduced lung capacity, and declining reproductive function^1–3^. These phenotypes reflect multi-scale remodeling within tissues, including shifts in cellular composition, extracellular matrix remodeling, and tissue-wide architectural integrity^1,4^. Despite its inevitability, the mechanisms driving aging and their consequences across tissues remain incompletely understood. Aging is associated with profound morphological and functional heterogeneity, both within and between tissues, presenting a challenge in linking cellular, microanatomical, and organ-level changes to organismal health.

Research on human aging has often focused on molecular, genomic, and cellular features, such as DNA methylation^5–7^, telomere attrition^8–10^ or cellular senescence^11,12^, offering valuable insights into processes that disrupt cellular homeostasis. While these molecular hallmarks of aging are increasingly well defined, their manifestation into spatially organized structural changes in tissue remains poorly understood. This limitation is compounded by the reliance on animal models, which do not fully recapitulate the complexity of human aging due to genetic, physiological, and environmental differences. Human-focused studies, particularly those leveraging large-scale repositories of data^13,14^, have nonetheless begun to address these gaps.

Human tissues are not homogeneous but are characterized by distinct microanatomical domains - recurrent cellular arrangements tailored to organ-specific functions^15–17^, making their detection a powerful lens for understanding tissue homeostasis and its dysregulation with age. For instance, the small intestine is critical for digestion and nutrient absorption, while simultaneously containing microbiota and defending against pathogens^18^. To perform these functions, it is divided into a layered structure of mucosa, submucosa, and muscularis of the gastrointestinal tract, which ensures nutrient absorption, barrier defense, and peristalsis^18^, respectively. While the compartmental organization of tissue is a long-recognized anatomical feature, we lack in its systematic and quantitative analysis, leaving the influence of aging on microanatomy across human tissues understudied. Understanding the impact of aging in these compartments is essential for uncovering universal tissue organization principles and prioritizing mechanistic investigations.

Recent advancements in computational pathology and deep learning^19,20^ have revolutionized the analysis of histopathological images, enabling a quantitative understanding of tissue structure on a scale that was not possible before. In our prior work, we developed a method for the unsupervised detection of tissue architecture using graphs (UTAG)^21^ in multiplexed imaging data. However, the limited field of view and high costs of multiplexed imaging preclude large sample sizes in the study of human aging, where large and diverse cohorts are essential to capture the substantial heterogeneity and multiple factors that influence the aging process.

Whole-slide hematoxylin and eosin (H&E) stained digital images on the other hand, are inexpensive and have a field of view that captures large tissue structures at single-cell resolution. Deep learning (DL) vision models excel at the extraction of rich morphological features in H&E images^20^ which can be used for downstream tasks such as cell detection and classification^22^, disease detection, classification^23–25^, and patient stratification^26–28^. This versatility enables the use of large (thousands of images) biobanks of histopathological images^29,30^ to systematically address how aging impacts tissue architecture across the human body^14^. At this scale, it becomes possible to build a comprehensive understanding of the heterogeneity inherent to the aging process by profiling thousands of individuals.

In this study, we introduce a systematic and quantitative investigation of aging at the mesoscale of human biology—the level of microanatomical domains that bridge cells and organs. To this end, we developed an unsupervised method to detect microanatomical domains on H&E images and applied it to the largest dataset of healthy human tissue: 40 tissues from 983 individuals across adulthood (20 to 70 years old). This approach enabled us to systematically characterize the identity, diversity, and dynamics of tissue organization with age across organs, establishing a quantitative framework to understand how aging and pathology reshape tissue-specific microanatomy across the human body.

## Results

### Unsupervised detection of microanatomical domains from histopathological images

To gain a systematic understanding of how aging reshapes the microscopic architecture of the human body (**Figure 1a**), we developed H&E-UTAG, a novel method for the unsupervised detection of microanatomical domains within H&E whole slide images (**Figure 1b, d**). Due to their wide field of view, histopathological images offer robust means to detect recurrent tissue organization patterns beyond local environments, and their spatial relationships in human organs (**Figure 1b**). The scalability of the method in combination with abundant images enables us to investigate age-related changes within individual domains across donors, as well as how aging alters the interplay between domains, ultimately impacting organ structure and function (**Figure 1c**).

**Figure 1.**
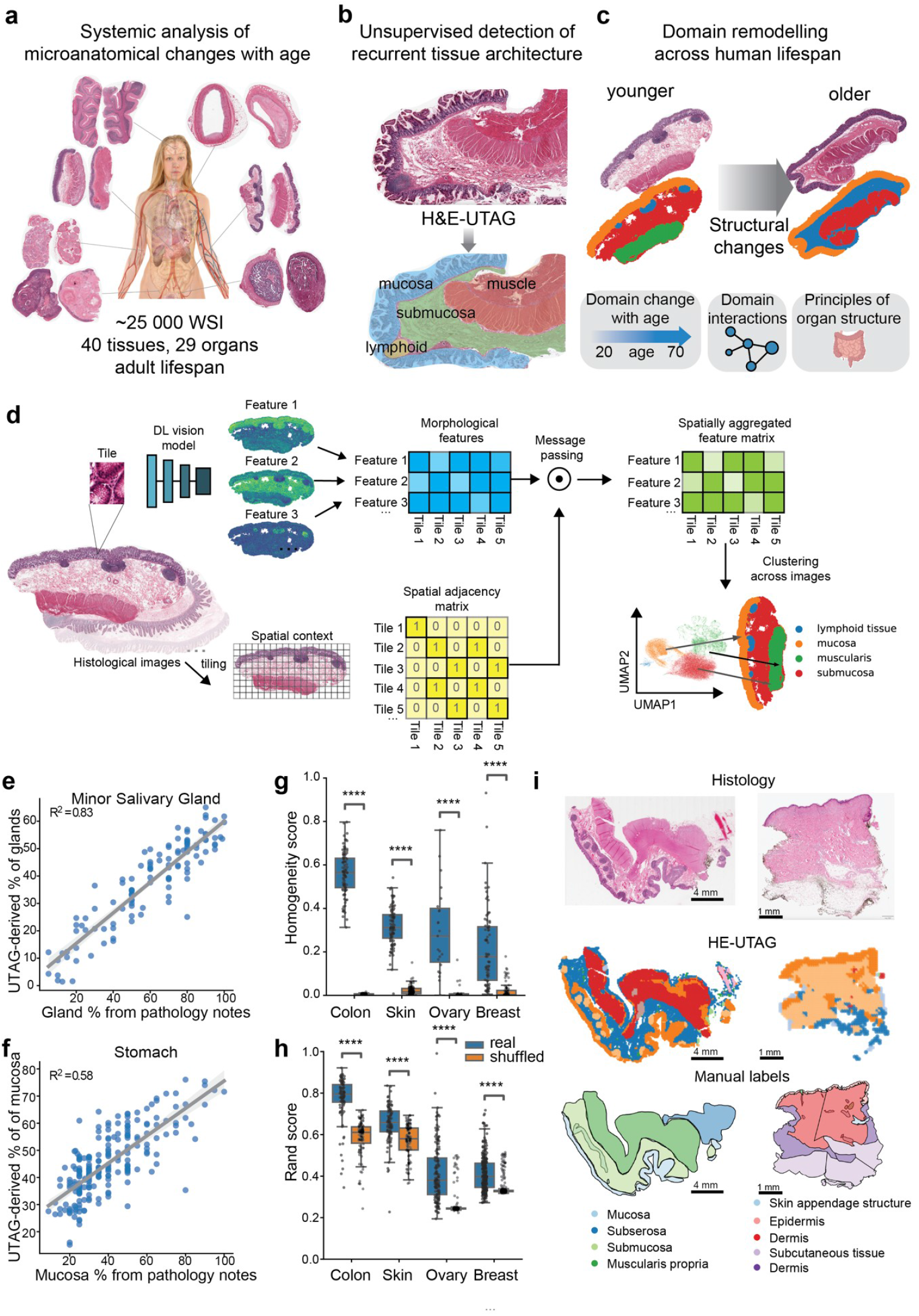
Unsupervised detection of microanatomical domains from histological images. **a**>, Study overview. The analysis encompassed 24 945 histological slides across 29 organs and 40 tissues from adult individuals across the lifespan that were used to systematically quantify age-associated microanatomical changes. **b**, Scheme of unsupervised detection of microanatomical domains with H&E-UTAG. **c**, Illustration of application of the H&E-UTAG to detect remodeling of microanatomy domains across age groups. **d**, Workflow of the H&E-UTAG pipeline. Histological images are tiled and passed through a deep learning vision model to extract morphological features. Spatial relationships between the morphological profiles of the tiles are encoded with message passing using morphological matrix and tile spatial adjacency matrix. To detect recurrent domains, unsupervised clustering is applied to the spatially aggregated feature spaces across images. **e, f**, Quantitative validation of H&E-UTAG detected minor salivary gland tissue and stomach mucosa with the percentage in the pathological notes. **g**, Homogeneity scores for H&E-UTAG derived clusters for colon, skin, ovary and breast in an external VSP dataset. **h**, Rand scores for H&E-UTAG derived clusters for colon, skin, ovary and breast in an external VSP dataset. **i**, Visual comparison of manual labels and H&E-UTAG derived labels in colon and skin from the VSP dataset.

H&E-UTAG is based on our previous work on multiplexed imaging^21^, but works on tessellated whole slide images, and leverages deep learning-derived features that capture morphological and textural properties of tissue. These features show distinct spatial distribution, often associated with specific biological structures (**Figure 1d**). By combining the features with their spatial relationships via message passing, the method propagates information, jointly encoding local morphology and surrounding tissue context **(Figure 1d)**. Clustering the integrated information across slides reveals recurrent archetypes of tissue organization, present within and across whole-slide images, enabling user-controlled tuning of granularity. H&E-UTAG is implemented in LazySlide^31^, which includes documentation and tutorials for its application.

To benchmark the method, we compared H&E-UTAG derived proportions of microanatomical domains with pathologist-annotated notes such as glandular tissue in salivary gland and stomach mucosa, and found strong correlations (**Figure 1e**, r^2^=0.83, r=0.91, p < 0.0001), (**Figure 1f**, r^2^ = 0.58, r=0.76, p < 0.0001), Pearson correlation). To assess generalizability, we applied H&E-UTAG to a dataset of 293 whole-slide images from colon (n = 98), skin (n = 96), ovary (n = 160) and breast (n = 291) with expert-annotated semantic segmentations^32^ (**Supplementary Figure 1a-b**). We assessed performance by comparing H&E-UTAG-derived domains to the manual ground truth using the Rand and homogeneity scores (see Methods). Across all tissues, H&E-UTAG output significantly outperformed randomized labels (p < 0.0001, **Figure 1g-h**), indicating that the method identifies spatially coherent and biologically meaningful microanatomical domains. Qualitative comparisons with expert annotations further confirmed high concordance across layered and morphologically complex tissues (**Figure 1i**).

### Microanatomical composition of the human body and its variation in aging and pathology

To investigate the microanatomical composition of the human body at scale, we applied H&E-UTAG to 24,945 whole slide images of healthy tissue from the GTEx project^33^ (162 million tiles; **Supplementary Figure 2a-b**). Across these 24,945 WSIs, we analyzed tissues from 971 individuals (age range 20-70 years, median 56 years; 66.8% male, 32.2% female; **Supplementary Figure 2c**,**e**), covering 40 tissue types from 29 organs (**Supplementary Figure 2d**,**e**), collected via a rapid autopsy protocol. To maximize comparability across organs, we used a pre-trained deep learning model for feature extraction, and the same tile size and clustering resolution across all images. This enabled us to jointly detect the microanatomical composition for hundreds of slides per organ without the need for manual labels. Across the dataset we identified 218 recurrent domain types, which we organized into 74 classes and 10 superclasses for interpretability (**Figure 2a, Supplementary Figure 3a-d**, Methods), collectively labeling more than 99% of the tissue area in the GTEx cohort (**Supplementary Figure 3b, Supplementary Figure 4**).

**Figure 2.**
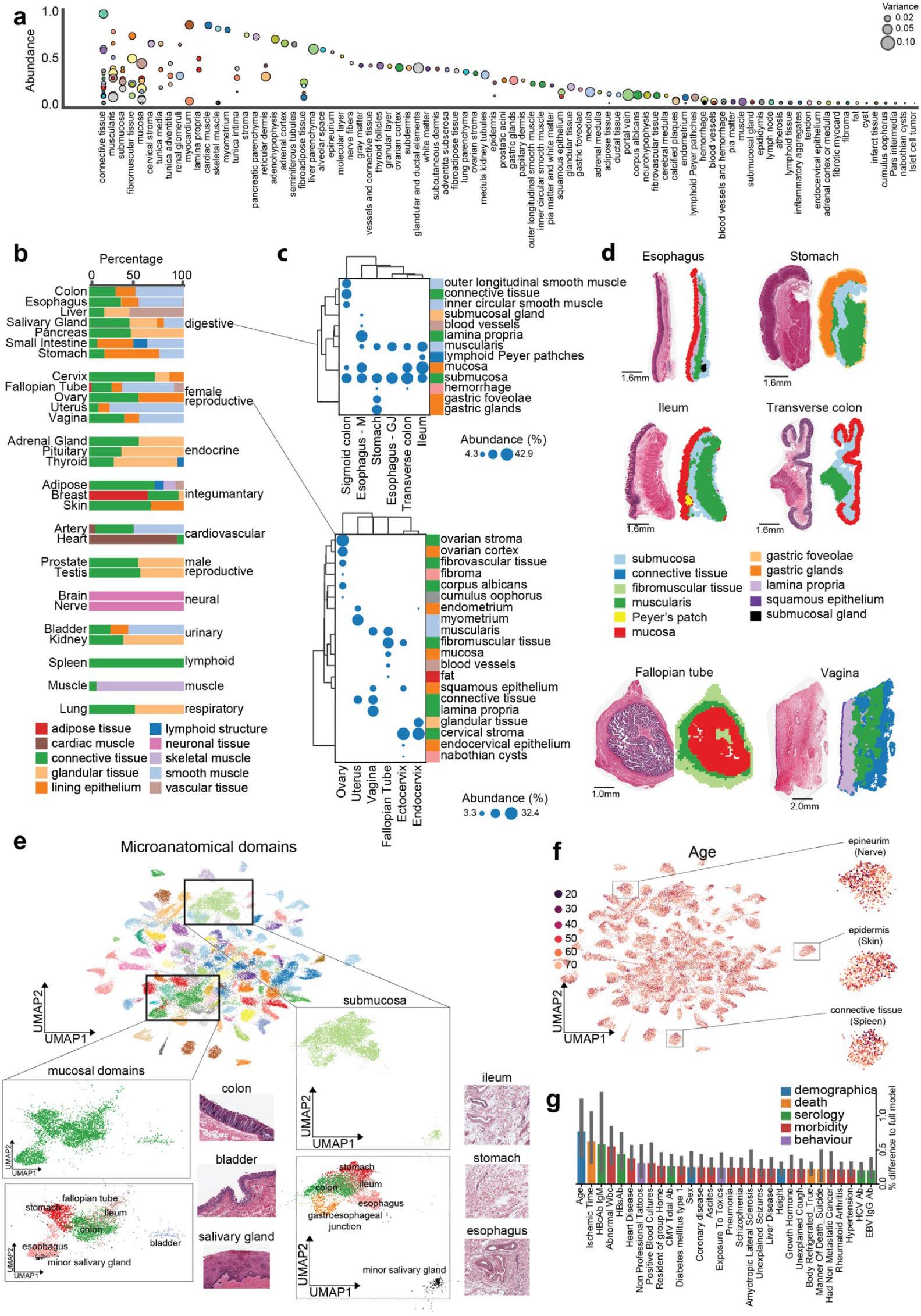
Microanatomical domain diversity, tissue composition, and age-associated remodeling across human organs. **a**>, Abundance and inter-individual variance of H&E-UTAG-derived microanatomical domains across all analyzed tissues in the GTEx dataset. Each dot represents a domain, colored by major tissue category, and sized by variance of abundance. **b**, Relative composition of tissue types across representative organs. **c**, Domain abundance profiles for tissues of the gastrointestinal tract (top) and female reproductive system (bottom), showing hierarchical clustering of domains (rows) and tissues (columns). **d**, Representative examples of unsupervised H&E-UTAG domain maps overlaid on histological sections of tissues from the gastrointestinal tract (esophagus, stomach, ileum, transverse colon) and female reproductive system (fallopian tube, vagina). **e**, UMAP embedding of all inferred microanatomical domains, illustrating morphological diversity. Insets highlight subsets of domains including mucosal and submucosal compartments, showing consistency of structure across distinct organs (e.g., colon, stomach, ileum, esophagus, salivary gland). **f**, UMAP embedding of all domain features colored by donor age, highlighting age-related spatial shifts in specific tissue types (e.g., connective tissue in the spleen, epidermis in the skin, epineurium in the nerve). **g**, The variance explained by metadata categories (demographic, death, serology, morbidity, behavior) for domain abundance across tissues.

Beyond changes in abundance, we also explored the morphological diversity of microanatomical domains by analyzing the deep learning-derived feature profiles for each domain in each tissue image (**Figure 2e**). We observed remarkable morphological diversity of microanatomical compartments across the human body. Some domain types, such as mucosal and submucosal compartments, grouped together across multiple organs, indicating that their morphology is more similar to each other than to other domains from the same organ (**Figure 2e**). Such cross-organ clustering suggests that these domains follow conserved structural “blueprints” that may be dictated by their specialized physiological roles, such as barrier formation or connective support. By contrast, other domain types such as connective tissue exhibited greater morphological heterogeneity between organs, potentially reflecting adaptation to local mechanical and functional demands (**Supplementary Figure 7a**). These patterns of conserved and variable domain morphology establish a framework to systematically map the factors reshaping universal and tissue-specific microanatomical organization across organs.

Using this framework, we next asked how these morphological features relate to chronological age of tissue donors. Appreciating how donors of different age distribute in the same unsupervised morphological embedding (**Figure 2f**), we observed age-associated gradients within certain domains, indicating progressive and measurable remodeling of their morphology over the adult lifespan. Given these morphological shifts, we utilized the domain abundance profiles to assess the contribution of age, and other donor characteristics, to overall variation in domain composition. We performed variance decomposition analysis using 111 annotated donor variables spanning demographic, clinical, serological, and technical categories (**Methods**). Across tissues, these factors together explained a substantial fraction of the variability in domain abundances (**Figure 2g**). Among them, donor age consistently accounted for the largest share of explained variance across organs, while other factors, such as organ-specific pathologies or comorbidities, contributed substantially to certain tissues. This highlights that, while age exerts a broad, cross-cutting influence on tissue composition, it acts alongside other factors that may dominate in specific tissues.

### Dynamics of functional domains across human lifespan

Because donor age emerged as the leading factor explaining the greatest variation in the abundance of microanatomical domains (**Figure 2g**), we next systematically quantified its influence across tissues.

To capture general trends, we restricted the analysis to microanatomical domains with abundance greater than 1% in each tissue, resulting in 128 domain-tissue pairs. Using generalized linear models, we estimated the relative change in domain abundance across the adult lifespan, adjusting for manner of death and post-mortem ischemic interval inherent to the cross-sectional GTEx cohort. We identified 37 out of 128 domains with significant age-associated changes in abundance (**Figure 3a, Supplementary Figure 8 and 9a-c**), with some domains increasing (**Figure 3d**) or decreasing (**Figure 3e**), with most changes being less than 3% per decade of adult life.

The magnitude and direction of microanatomical changes varied across organ systems and tissue types. The integumentary and reproductive systems were among the most variable, whereas the nervous and muscular systems remained comparatively stable (**Figure 3b**). At the level of broad tissue classes, connective, glandular, and epithelial tissues were most affected, while skeletal muscle and lymphoid structures were largely spared from age-associated abundance shifts captured by histology (**Figure 3c**).

The strongest change detected was an increase in corpus albicans - the scar that accumulates with ovulation cycles^34^. Its proportion in the ovary increased from a mean of 1.4% in 20-year-olds, to 11.89% in 70-year-olds, a 3.51% increase per decade (**Figure 3g**). We also observed changes in microanatomical composition which had not been previously reported. The submucosal layer of various gastrointestinal tract tissues declined from 42.9% to 34.5% between ages 20 to 70 (-0.98% per decade) (**Figure 3f**). The adenohypophysis domain of the pituitary gland decreased from 82.6% to 67.9% (-3.42% per decade) (**Figure 3h**), a trend previously noted^35,36^, but of unclear physiological relevance. Additionally, we observed an age-associated increase in connective tissue in many organs (**Figure 3h, Supplementary Figure 8b**). Collectively, these findings underscore that even at the microanatomical level, tissue composition undergoes measurable and tissue-specific remodeling throughout adulthood.

**Figure 3.**
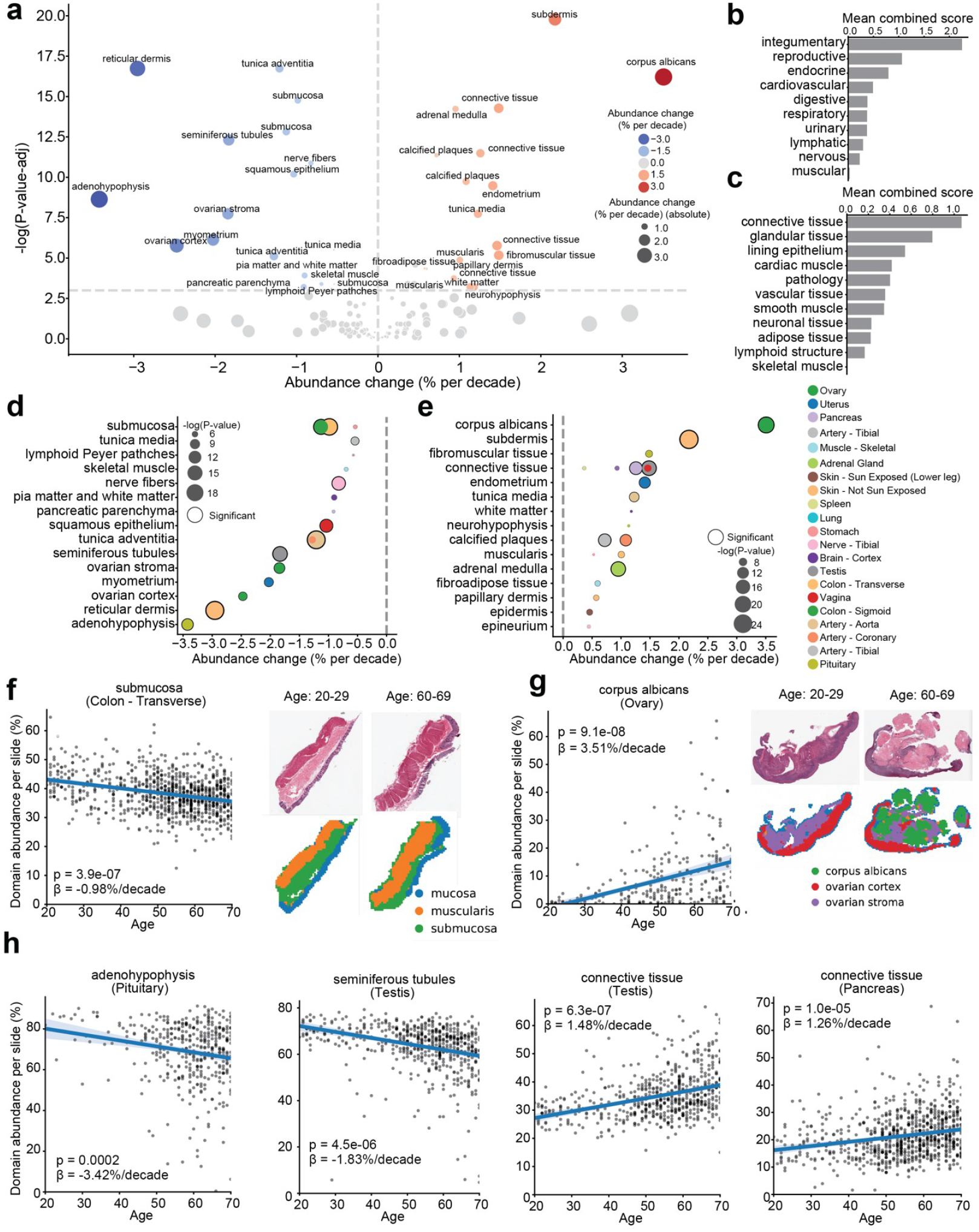
Age associated remodeling of the microanatomy. **a**>, Volcano plot of microanatomical domains abundance change with age. **b**, Mean combined score of microanatomical age change across organ systems. **c**, Mean combined score of microanatomical age change across tissue types. **d**, Microanatomical domains with decreased abundance with age **e**, microanatomical domains with increased abundance with age. **f**, Abundance change in submucosa with age with a visual example. **g**, Abundance change of corpus albicans with age with a visual example. **h**, Abundance change of adenohypophysis, seminiferous tubules, testis connective tissue and pancreatic connective tissue with age.

### Subclinical pathology drives selective microanatomical remodeling across human tissues

Aging is closely associated with progressive tissue dysfunction and chronic disease, yet its earliest manifestations often starts subclinical. To investigate if and how such early pathology influences tissue microanatomy, we analyzed the existing systematic annotation of pathological features in the GTEx cohort (9643 instances spanning 54 types). We distinguished between two complementary dimensions of microanatomical remodeling: domain abundance, reflecting shifts in tissue composition, and domain morphology, capturing subtler alterations such as cellular reorganization or extracellular matrix remodeling. This framework (**Figure 4a, Supplementary Figure 10**) allowed us to systematically assess whether pathology primarily affects the amount of a structural compartment, its intrinsic characteristics, or both.

We identified multiple domain types that exhibited significant alterations in either abundance (**Figure 4b, Supplementary Figure 10a**) or morphology (**Figure 4b, Supplementary Figure 10b-d**) in the presence of pathology. Not all microanatomical domains within a given tissue were equally affected, indicating variable susceptibility and resilience across domain types. Pathologies also differed in their predominant mode of impact: some primarily altered domain abundance, while others affected morphology. For instance, testicular atrophy was associated with morphological changes in the seminiferous tubules (**Figure 4c**), likely due to the previously described decrease in spermatogenesis with age^37,38^. We also observed an increase in connective tissue with atrophy, replacing the area occupied by seminiferous tubules (**Figure 4c**). In contrast, arterial fibrosis increased the abundance of the tunica adventitia but produced minimal morphological changes (**Figure 4c**). Finally, liver cirrhosis was characterized by both expansion of connective tissue and marked morphological remodeling (**Figure 4c**).

**Figure 4.**
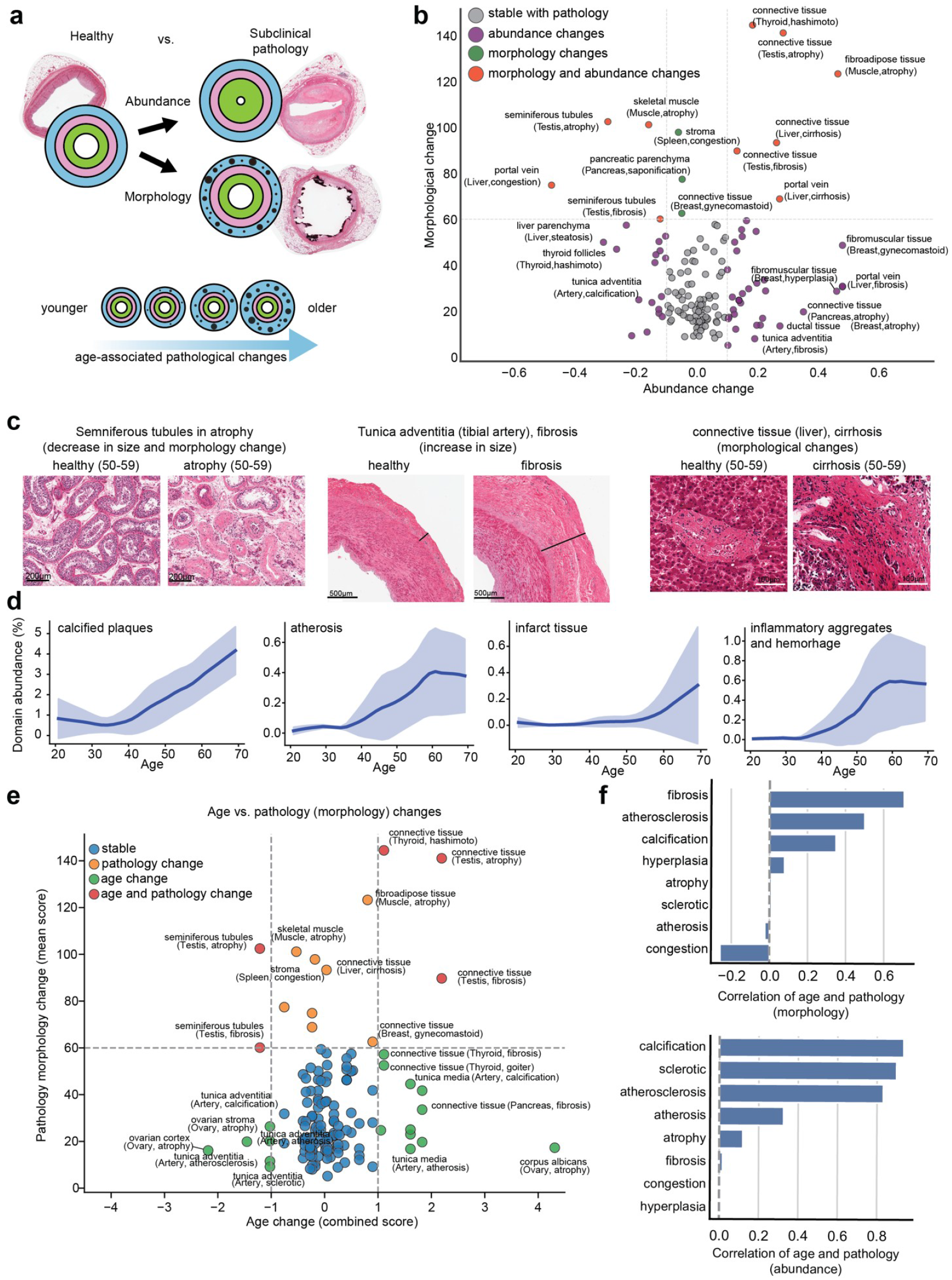
Effects of subclinical pathology on the microanatomy of human tissues. **a**>, Scheme of the effects of subclinical pathology on the abundance and morphology of microanatomical domains **b**, Abundance and morphological changes in the microanatomical composition associated with subclinical pathologies. **c**, Visual examples of pathology-related changes of seminiferous tubules, tunica adventitia in the tibial artery and liver connective tissue. **d**, Age-dependent accumulation of pathology-associated domains in calcified plaques, atherosclerosis, infarct tissue, and inflammatory aggregates. **e**, Comparison of pathology-driven and age-driven microanatomical changes. **f**, Correlations between age and pathology domain changes for specific pathology classes. Top: correlation of age with pathology-associated morphological change; Bottom: correlation of age with pathology-associated domain abundance. Fibrosis and atherosclerosis show the strongest age-related correlation across both metrics.

At the superclass level, organs varied widely in the proportion of epithelial, connective, muscle, and other tissue types (**Figure 2b, Supplementary Figure 5a, Supplementary Figure 6**), providing a comparative baseline from which to explore the recurrence or restriction of specific microanatomical domains across the body. Across organ systems, microanatomical composition reflected both shared architectural motifs and highly specialized domains (**Figure 2b-d**). The gastrointestinal tract displayed a conserved tripartite layering of mucosa, submucosa, and smooth muscle across segments, reflecting its roles in nutrient absorption, barrier defense, and peristalsis (**Figure 2c-d**). In contrast, the female reproductive system combined recurrent domains such as mucosa and muscularis with specialized ones, including the ovary-restricted cumulus oophorus, which encapsulates and nourishes the oocyte (**Figure 2c-d**). These examples illustrate how recurrent domains support conserved physiological functions across anatomically distant sites, while unique domains underpin organ-specific roles.

For ten pathological instances, including arterial calcification, H&E-UTAG identified domains that primarily represent the pathology within the tissue. Leveraging these, we explored how the abundance of these pathology-associated domains changes with age (**Figure 4d**). Arterial calcified plaques increased gradually with age, with an accelerated rise after 40 years of age. Conversely, atherosis and cardiac inflammatory aggregates peaked around 60 years of age, while infarct tissue remained scarce until it sharply increased after 60 years (**Figure 4d**).

Age and age-associated pathology are often conflated, with structural changes in older tissues implicitly attributed to disease processes. To test whether healthy aging remodels the same microanatomical domains as pathology, we compared the strength of age-abundance associations in non-pathological domains (from **Figure 3**) with the magnitude of pathology-morphology associations in the same tissues (from **Figure 4b**) (**Figure 4e**). The relationship was generally weak: domains most affected by healthy aging were not necessarily those most altered in the presence of pathology. Only a minority of domains showed strong effects in both contexts, suggesting that normal aging and pathology frequently remodel different structural components of tissue indicating they follow partially distinct microanatomical trajectories.

We next asked whether specific pathologies recapitulate the same microanatomical shifts observed with aging. Correlating pathology-associated changes in abundance and morphology with age-related changes across domains revealed that fibrosis, atherosclerosis, and calcification showed the strongest positive correlations in morphology, whereas vascular congestion exhibited a weak negative correlation (**Figure 4f**, top). In terms of abundance, coronary pathologies such as calcification, sclerosis, and atherosclerosis aligned most closely with age-related changes (**Figure 4f**, bottom).

These results show that tissue-specific pathology is associated with selective remodeling of tissue architecture, impacting both abundance and morphology of microanatomical domains to varying degrees.

### Aging alters higher-order spatial interactions between tissue domains

Microanatomical domains are fundamental units of organ architecture, and their coordinated arrangement collectively underpins tissue function. Using our systematic and quantitative atlas of human microanatomy, we built a multi-scale hierarchical representation of organ architecture, capturing both anatomical subdivisions within the same organ (e.g., cerebellum vs cortex in the brain, or transverse vs sigmoid colon) and the classes and subtypes of microanatomical domains (e.g., epithelium, adipose, vascular, muscle tissue; including specific subtypes such as mucosal or squamous epithelium) (**Figure 5a**). This framework reveals both the shared and unique domain composition across organs, highlighting how conserved microanatomical features are deployed in tissue-specific contexts across organs.

**Figure 5.**
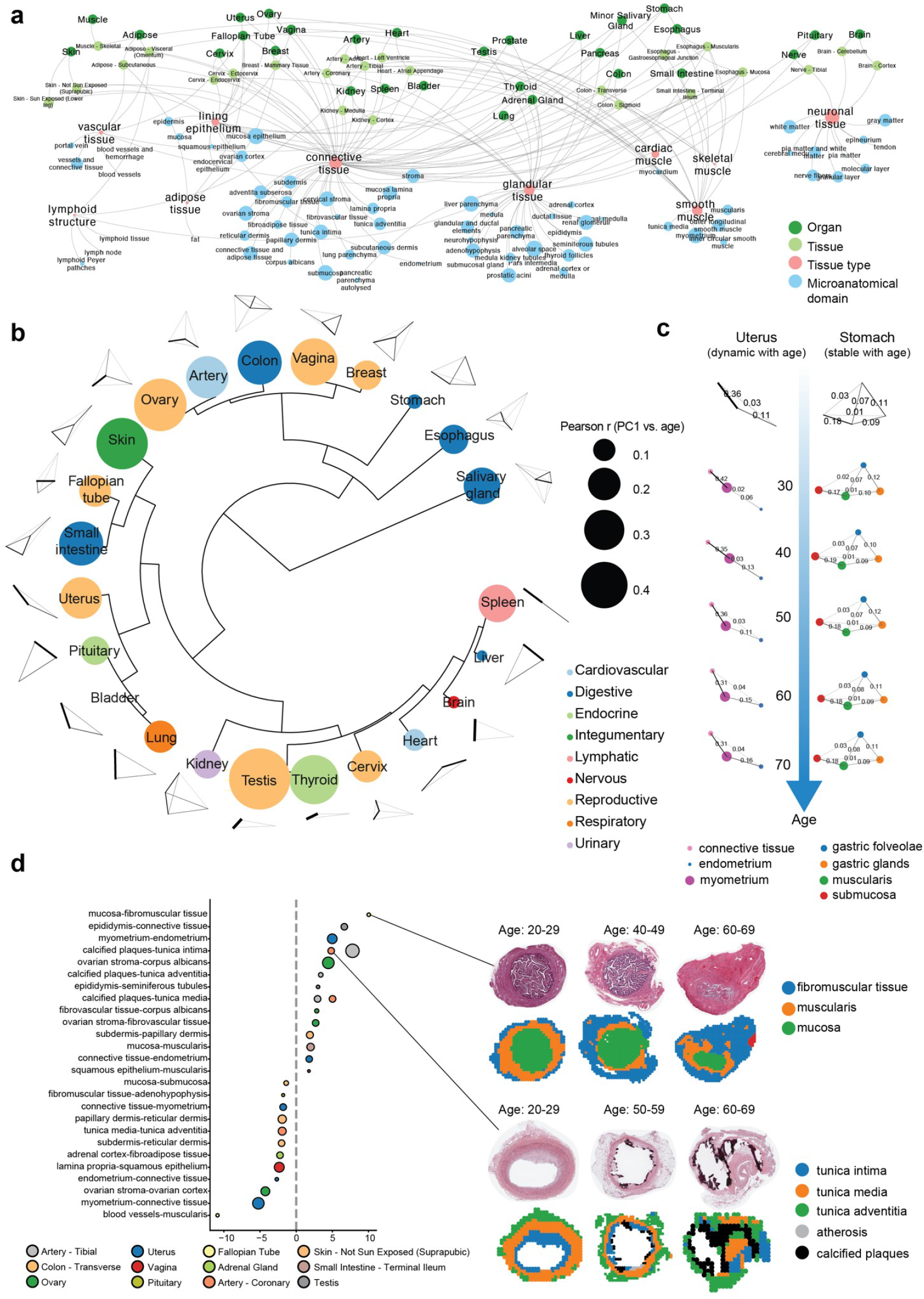
Domain interaction networks and dynamic remodeling with age across tissues. **a**>, Hierarchical representation of organ, tissue, tissue type, and microanatomy organization in the human body. Nodes represent individual organs, tissues, tissue types, and microanatomical domains; edges indicate hierarchical associations across anatomical scales **b**, Hierarchical clustering of organs based on the structural properties of their domain interaction networks. Representative interaction graphs are shown below each organ. The node color represents organ system and node size represents Pearson r correlation of the PC1 of the interaction embedding and age. **c**, Age-stratified domain interaction networks in two representative organs: stomach (stable across age) and uterus (dynamic across age). **d**, Quantification of age-dependent changes in domain-domain interactions across tissues with representative histology and H&E-UTAG segmentation examples of the fallopian tube and coronary artery across selected decades of life.

We used this atlas to examine how aging influences the organization of tissue architecture at a higher-order level. By quantifying physical contacts between microanatomical domains, we captured patterns of domain interaction across organs (**Supplementary Figure 11a**, Methods). This allowed us to cluster tissues based on their global architectural properties and identify distinct architectural archetypes (**Figure 5b, Supplementary Figure 11b**). To quantify the effect of age on each tissue’s global architecture, we used topological features of their structure to predict the age of each donor and quantified the amount of topological architecture explained by age (**Supplementary Figure 11c-e**). This highlighted tissues where the dominant axis of structural variation is age-associated or stable (**Supplementary Figures 12-13**). Organs such as the uterus showed pronounced age-associated remodeling, where the endometrium is more strongly associated with connective tissue in older age (**Figure 5c**), but in most organs the overall topology remained remarkably resilient despite pervasive changes in abundance, morphology and pathology (Figures 3-4).

Finally, given that organ function emerges from the coordinated interactions of microanatomical domains, we examined how direct physical interactions between domains change with age, modeling their pairwise interactions across organs (**Figure 5d, Supplementary Figure 11e-f**). Across tissues, we identified widespread remodeling of cross-domain interactions with age, with some interfaces increased in contact while others progressively lost connectivity. Notably, the proportion of mucosa-fibromuscular interactions in the ovary and contacts between calcified plaques and the tunica intima in coronary and tibial arteries increased with age (**Figure 5d**), whereas subdermis-reticular interactions in skin and muscularis-submucosa interaction in colon declined significantly with age (**Figure 5d**). The fallopian tube showed pronounced remodeling, with reduced mucosa-muscularis and increased mucosa-fibromuscular interactions (**Figure 5d**), suggesting an age-related replacement of contractile tissue essential for oocyte transport and consistent with reproductive decline. In arteries, the increased contact between calcified plaques and the tunica intima points to structural shifts linked to pathological calcium deposition.

Overall, aging selectively reshapes local interactions between microanatomical domains in a tissue-specific manner, subtly altering organ architecture but largely preserving global structural organization. These changes likely contribute to gradual functional decline and highlight the importance of understanding tissue remodeling at multiple spatial scales.

## Discussion

In this study we described the microanatomical composition of the human body and systematically quantified how it changes with age and pathology. Specifically, we developed a novel unsupervised method to detect the microanatomy from whole slide H&E images and have applied it to the largest collection of healthy human tissue samples to date. This analysis revealed the remarkable morphological diversity and organizational complexity of human organs. Age emerged as the strongest factor influencing the abundance of microanatomical domains, with 28% showing significant age-associated changes. Despite these shifts, the higher-order architecture of organs remains largely resilient. By focusing exclusively on human tissue, our work captures the complexity and heterogeneity of human aging, offering insights that cannot be fully captured in animal models.

Tissue-specific factors, such as extracellular matrix remodeling are closely linked to the pathophysiological hallmarks of aging^4,39–41^. In our previous work we demonstrated that aging strongly influences tissue morphology, enabling the prediction of biological age from histopathological images^14^. However, the specific tissue elements underlying these changes were not well defined. The present study addresses this gap by detecting and quantifying the microanatomy of human tissue in a systematic way and quantitatively associating it with age. In some organs, including the female and male reproductive system, we observed significant microanatomical shifts with age, while some other domains such as the cerebellar medulla remained remarkably resilient to both age and pathology-related alterations.

Age-associated remodeling of discrete microanatomical domains across the human body aligns with multiple cellular and molecular hallmarks of aging. The increase in connective tissue across diverse organs is consistent with extracellular matrix remodeling and fibrosis, hallmarks of altered intercellular communication and chronic low-grade inflammation^42–44^. The progressive loss of functional compartments such as seminiferous tubules in the testis, adenohypophysis in the pituitary, and the gastrointestinal submucosa likely results from reduced regenerative capacity^45,46^ and disruption of the stromal and vascular niches that maintain tissue homeostasis^47^. Differential susceptibility to age-related microanatomical changes may have evolutionary roots: tissues in the nervous and muscular systems show remarkable domain composition stability, whereas the integumentary and reproductive tissues are highly plastic, potentially reflecting an evolutionary prioritization of barrier defense and reproductive capacity during the reproductive lifespan, in line with theoretical predictions^48^.

A key advance of this work is the ability to computationally separate age-associated changes in histologically normal tissue from those driven by subclinical pathology. This distinction is rarely achievable in large-scale human histology datasets due to the use of clinically sourced samples, but it is essential for disentangling the biology of physiological aging from early disease. We find that domains most affected by healthy aging are often distinct from those most remodeled in pathological states, suggesting partially diverging structural trajectories. For example, ovarian corpus albicans accumulation is a hallmark of reproductive aging independent of ovarian pathology, whereas arterial calcification and fibrotic replacement of myocardial tissue follow a pathology-aligned trajectory that correlates with age but are mechanistically distinct. These findings challenge the assumption that structural changes in older tissues are inherently disease-driven and provide a framework for defining domain-level biomarkers of healthy versus pathological aging.

While the GTEx cohort is an outstanding resource for studying tissue microanatomy, it has important limitations. The donor population is male-biased (66%), and organ coverage is uneven (881 samples for the most represented tissue, 37 in the least). The data are cross-sectional and post-mortem, preventing longitudinal follow-up and the introduction of potential artifacts from ischemic time or cause of death cannot be completely excluded, despite computational modeling using excellent annotation of technical confounders. Lifestyle and environmental exposures are largely unrecorded, limiting assessment of extrinsic drivers of microanatomical change, despite the extensive and valuable information on demographic and clinical variables in the cohort. Additionally, the absence of health outcomes makes it difficult to directly link structural remodeling to physiological decline. Future studies that integrate spatially resolved molecular assays with longitudinal sampling and functional phenotyping will be critical to fully understand how extrinsic factors shape human tissue aging.

In summary, our study introduces a new and comprehensive view of human tissue microanatomy, revealing fundamental principles of organ organization and how these structures are altered during aging and disease. By systematically mapping microanatomical domains across the body, we show how age and pathology differentially remodel tissue architecture, highlighting both resilient and labile microanatomical domains and their interactions. This framework provides a crucial foundation for understanding the mechanisms of tissue aging, the development of age-related diseases, and for designing diagnostic and therapeutic strategies targeting tissue-specific pathology.

## Methods

### H&E-UTAG

This method is based on UTAG, a previously developed method for the unsupervised discovery of tissue anatomy with graphs^21^. The input to the method is a morphological matrix and a spatial adjacency matrix for a number of whole slide images. To derive the morphological matrix for each image, we used features extracted from a AlexNet model^49^ pretrained on ImageNet^50^ for tiles of the whole slide image. Specifically, non-overlapping tiles of size ∼112μm × 112μm (224 pixels at 20X magnification) were extracted per image for feature generation. Features were extracted from the last linear layer and resulted in a feature vector of shape 1000 per tile. To get the spatially aggregated features, we performed matrix multiplication of the morphological matrix (N morphological features X M tiles) and the l1-normalised spatial adjacency matrix of the tiles (M tiles X M tiles) for each histopathological image separately. To detect microanatomical domains, we clustered all tiles from all images of the same tissue separately with Leiden clustering^51^ with resolution 0.4 fixed for all tissues. After clustering, cluster labels were assigned back to their location in the tissue. For interpretability, we manually assigned hierarchical labels to the anatomical clusters by visual inspection of the clusters.

### Benchmarking with semantic labels

To benchmark the detection of microanatomical domains on fully annotated whole slide images, we leveraged the Visual Sweden Project datasets for breast, colon, skin and ovary^32^, which contain semantic labels generated by experts. We applied H&E-UTAG as described above to the entire set of the whole slide images (n=740), separately for each tissue without restricting clustering to annotated regions, to preserve the unsupervised nature of the method. For a quantitative evaluation, we focused on the slides with more than one annotation (n=296, Colon=98, Skin=96, Breast=79, Ovary=23), because slides with single annotation (e.g. only large tumor region annotated) produced trivial binary scores and were not informative for benchmarking. For each tile we determined the corresponding pathologist-assigned manual annotations. To avoid ambiguity, we excluded tiles with overlapping labels (e.g., a mucosal region embedded within submucosa with both “mucosa” and “submucosa” labels). We also excluded unlabeled tissue pieces when only part of the slide was annotated. The final benchmarking set therefore consisted only of tiles with a single, unique label assignment. On this set, we calculated homogeneity and Rand scores using scikit-learn^52^ for each slide separately.

### Benchmarking of slide-level labels

We assess how well inferred H&E-UTAG domains reflect expert annotated features at slide level by utilizing the pathological notes from the GTEx cohort that had annotated domain proportions in minor salivary gland and stomach. First, we parse quantitative proportional information from natural language notes when available. Only slides with available information were considered in this analysis. These contained information such as: “2 pieces, ∼10% is salivary gland elements with mild chronic inflammation, the remainder is lip, stroma, skeletal muscle”. When tissue pieces exhibited varying proportions of the domain of interest (e.g., “2 pieces of glandular elements (delineated) are ∼60-70% of tissue, rest is stroma/squamous mucosa”), the average was used. Cases lacking the domain of interest (e.g. “2 pieces; lip and skeletal muscle, no glands”) were annotated as 0% and included in the averaging process when applicable. Finally, we manually reviewed all extracted values to correct any inconsistencies. We then compared the annotated proportions with H&E-UTAG-derived by calculating Pearson correlation and r-square value with r2_score() function from scikit-learn^52^ package.

### Morphological profiling of the microanatomical domains and UMAP construction

To construct a UMAP of morphological features across domains, we mean aggregated the morphological features for each tile in a single image by the microanatomical domain. We applied Principal Component Analysis (PCA) and derived 50 principal components from the mean morphological features, followed by uniform manifold approximation and projection (UMAP) from the Scanpy^53^ package.

### Variance explained analysis

Analysis of variance was performed as described previously^14^. To quantify how biological and technical factors contribute to the variance in microanatomical domain abundance, we used a linear modeling approach based on the PCA of the matrix of samples and domain abundances. For each GTEx tissue separately, we computed principal components (PCs) and retained PCs that explained >1% of the variance. We then fit regression models using the statsmodels^54^ package to estimate how well a set of donor-level covariates could predict each PC. Covariates included age, sex, body mass index (BMI), cause of death, ischemic time, tissue procurement variables, and comorbidities (e.g., diabetes, dialysis), drawn from the GTEx metadata (see below). Categorical variables were one-hot encoded, and variables with no variance or high missing values were excluded. To assess the individual contribution of each covariate, we used a leave-one-out approach: for each PC and covariate, we fit a full model with all covariates and compared it to a reduced model excluding the covariate of interest. The resulting drop in R^2^ (or adjusted R^2^) quantified the unique explanatory power of that covariate. To compute an overall contribution per covariate, we scaled the R^2^ difference for each PC by that PC’s explained variance and summed across PCs. All models were fit independently per tissue, and results were aggregated to yield tissue-level estimates of factor importance in explaining variation in microanatomical domain abundances.

### Changes in microanatomical domains with age

To quantify age-related changes in the abundance of microanatomical domains, we systematically analyzed their prevalence across our dataset. Regression models (statsmodels^54^ implementation) were fitted for each domain to evaluate the relationship between domain abundance and age, while adjusting for potential confounders, including ischemic time, Hardy scale, and cohort annotations. Statistical significance was assessed using adjusted p-values with the Benjamini-Hochberg procedure for false discovery rate (FDR) correction (statsmodels^54^ implementation) and effect sizes were expressed as changes in domain abundance per decade of age.

### Changes in microanatomical domains with subclinical pathology

We separately quantified morphological changes, and changes in the abundance of the microanatomical domains. To quantify the changes in the abundance between healthy and pathology, we have trained a generalized linear model (GLM) with negative binomial distribution implemented in the statsmodels^54^ package. For each tissue and pathology pair, we modeled the number of tiles assigned to a specific domain as a function of pathology status (binary). Age (centered and scaled) was included as a covariate to control for age-related variation. We also included the total number of tiles per slide (centered and scaled) to account for variation in tissue area. This approach allowed us to estimate the pathology-associated change in domain abundance while adjusting for both age and sample size.

To quantify the morphological features between healthy and pathological tissues, we extracted the features using the visual-language model PLIP^55^ for all tiles in each image. For each microanatomical domain in each tissue, we then compared the average morphological profile between healthy and pathological samples. To account for differences in sample size between groups, we randomly subsampled the larger group to match the smaller one. We then used the rank_genes_groups() function from the Scanpy package^53^ to compute a score for each morphological feature that integrates effect size and significance. Finally, we summarized the overall morphological divergence between groups by taking the average difference score across features for each domain.

10 pathological domains that were detected as stand-alone domains are: ‘calcified plaques’, ‘atherosis’, ‘hemorrhage’, ‘nabothian cysts’, ‘inflammatory aggregates and hemorhage’, ‘fibrotic myocard’, ‘infarct tissue’, ‘fibroma’, ‘Islet cell tumor’, ‘cyst’.

### Comparison of changes with pathology and age

To compare changes with pathology and age, we compared morphological changes observed in the pathological analysis with the age-associated abundance changes from Figure 3a. To combine effect size and significance of age-associated change rate, we multiplied the log-fold change and -log10 of adjusted p-value (from Figure 3a) to get a combined score on Figure 4e (x-axis). The morphological score was derived in the same manner.

### Construction of domain interaction graphs

To represent each slide’s tissue architecture as a graph, we built graphs where microanatomical domain are nodes. For each domain, we identified its boundary tiles, defined as tiles that are adjacent to any other tissue tile (excluding lumen or empty space). We then quantified edges between domains by counting, for each pair of domains within a slide, the number (or proportion) of boundary tiles from the first domain that directly contact tiles of the second domain. These directed counts were used as edge weights in the resulting adjacency graph, thereby capturing the local physical interface between domains. Directed edges captured spatial relationships between domains, where edge weights represented the proportion of shared boundaries (i.e., touching tiles) between domain pairs. These proportions were computed for every domain pair within a given tissue. To assess age-associated remodeling of domain interactions, we fitted regression models for each domain pair, using interaction strength as the predictor and donor age as the outcome. The resulting regression slope quantified the effect size, and the associated p-value was used to assess statistical significance.

### Quantifying age-associated structure in domain interaction embeddings

To assess whether domain interaction patterns contained information related to chronological age, we analyzed interaction adjacency matrices using principal component analysis (PCA). For each tissue, we constructed a subject-by-feature matrix where features corresponded to pairwise domain interaction counts between anatomical domains. These matrices were centered and scaled prior to dimensionality reduction. PCA was then applied independently within each tissue, and the resulting principal component (PC) coordinates were used as embeddings for downstream analyses. We quantified the relationship between PCA embeddings and age using two complementary approaches: i) continuous correlation with age and ii) separation of age extremes. For each tissue, we computed the Pearson correlation coefficient between chronological age and the first principal component (PC1), which captures the dominant axis of variance across samples. The absolute value of the correlation coefficient was used as a measure of age-association, and statistical significance was assessed by permutation testing (randomly shuffling subject ages 100 times and recomputing correlations) (Supplementary Figure 11d). Furthermore, to test whether the embeddings capture separable structure in younger versus older individuals, we stratified samples into quintiles of age and retained the youngest (Q1) and oldest (Q5) groups. Silhouette coefficients were then calculated using these two groups as class labels and the full set of PCA embeddings as input. The silhouette score summarizes how well samples cluster by age extreme relative to their within-group similarity. As with the correlation analysis, a permutation baseline was established by randomly shuffling age labels 100 times and recomputing silhouette scores (Supplementary Figure 11d). The dot size on the circular clustermap on Figure 5b is the silhouette score.

### Topological profiling of tissue graphs

We computed topological metrics for the domain interaction graphs (section above) using scikit-network^56^. For each slide, we calculated a set of topological descriptors including basic graph descriptors (number of nodes, edges, connected components, largest connected component size, number of cycles, number of triangles) and structural signatures (Weisfeiler-Lehman colors, clustering coefficient, clique counts up to the graph degeneracy bound, mean PageRank, Katz centrality, HITS, betweenness centrality, closeness centrality, and degree) implemented in the scikit-network package^56^. Metrics were calculated for each slide and aggregated within each tissue by both mean and standard deviation. These aggregated metrics were centered and scaled and clustered hierarchically across tissues using the Pearson correlation distance and average linkage. For visualization, circular heatmap was done with Euclidian distance and unweighted pair group method with arithmetic mean (UPGMA) linkage with the R package ggtree^57^.

## Data availability

GTEx whole slide images are available from its portal (https://gtexportal.org).

## Code availability

Source code is provided as Supplementary Information files for reviewers and will be publicly available at the Github repository https://github.com/RendeiroLab/human_microanatomy after peer review.

## Acknowledgments

This project has received funding from the European Research Council (ERC) under the European Union’s Horizon Europe research and innovation programme (grant agreement no. 101220825). This research was funded by Angelini Ventures S.p.A. Rome, Italy. The Genotype-Tissue Expression (GTEx) Project was supported by the Common Fund of the Office of the Director of the National Institutes of Health, and by NCI, NHGRI, NHLBI, NIDA, NIMH, and NINDS. The datasets used for the analyses described in this manuscript were obtained from dbGaP at http://www.ncbi.nlm.nih.gov/gap through dbGaP accession number phs000424.v9.p2.c1. We thank Junbum Kim and the members of the Rendeiro group for their comments on the manuscript.

