## Supplementary Figures for "Systematic mapping of human tissue microanatomy reveals age-associated remodeling and resilience"

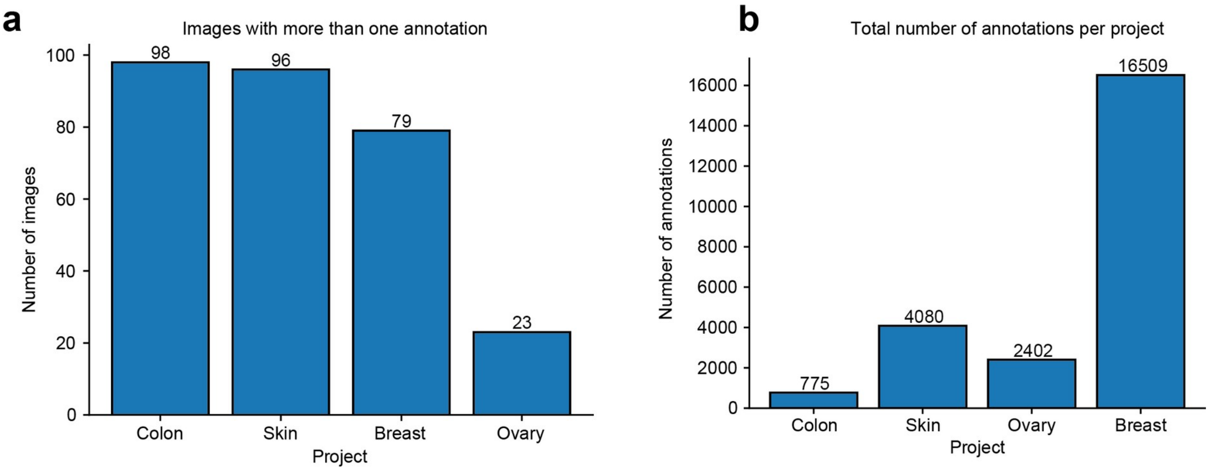


**Supplementary Figure 1. Overview of Visual Sweden Project dataset. a,** Cohort size per tissue for Visual Sweden Project used to benchmark H&E-UTAG (Colon, Skin, Ovary, Breast), only images with two or more pathologist annotations were used. **b,** Total number of expert annotations per cohort.

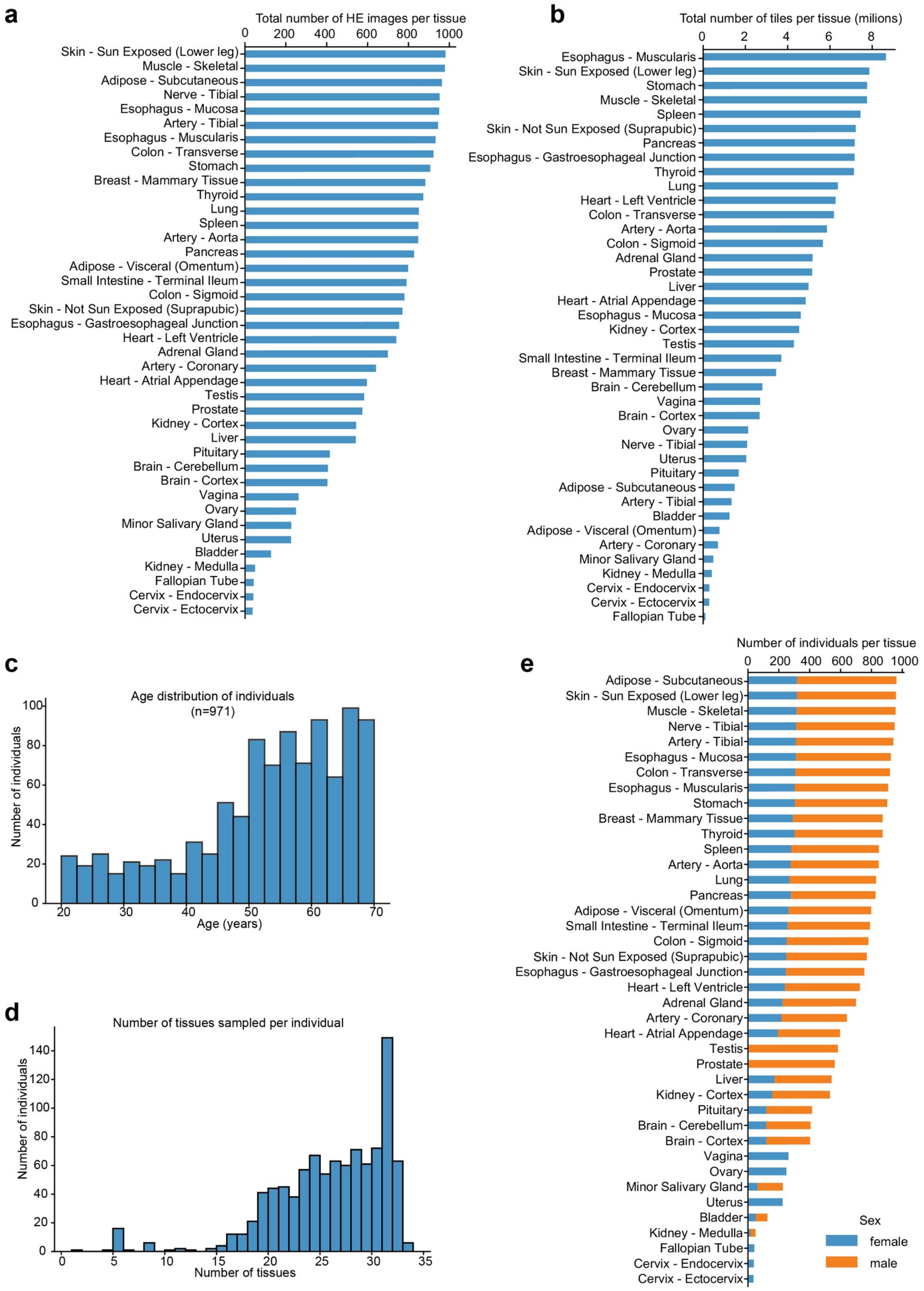


**Supplementary Figure 2. Overview of the GTEx dataset. a,** Total number of hematoxylin and eosin (HE) stained images per tissue type in the GTEx cohort. **b,** Total number of image tiles (in millions) per tissue. Tiles are non-overlapping image crops used for computational feature extraction. **c,** Age distribution of donors in the GTEx dataset (n=971 individuals). **d,** Distribution of the number of tissues sampled per individual, demonstrating comprehensive tissue sampling per donor. **e,** Number of individuals per tissue, colored by sex.

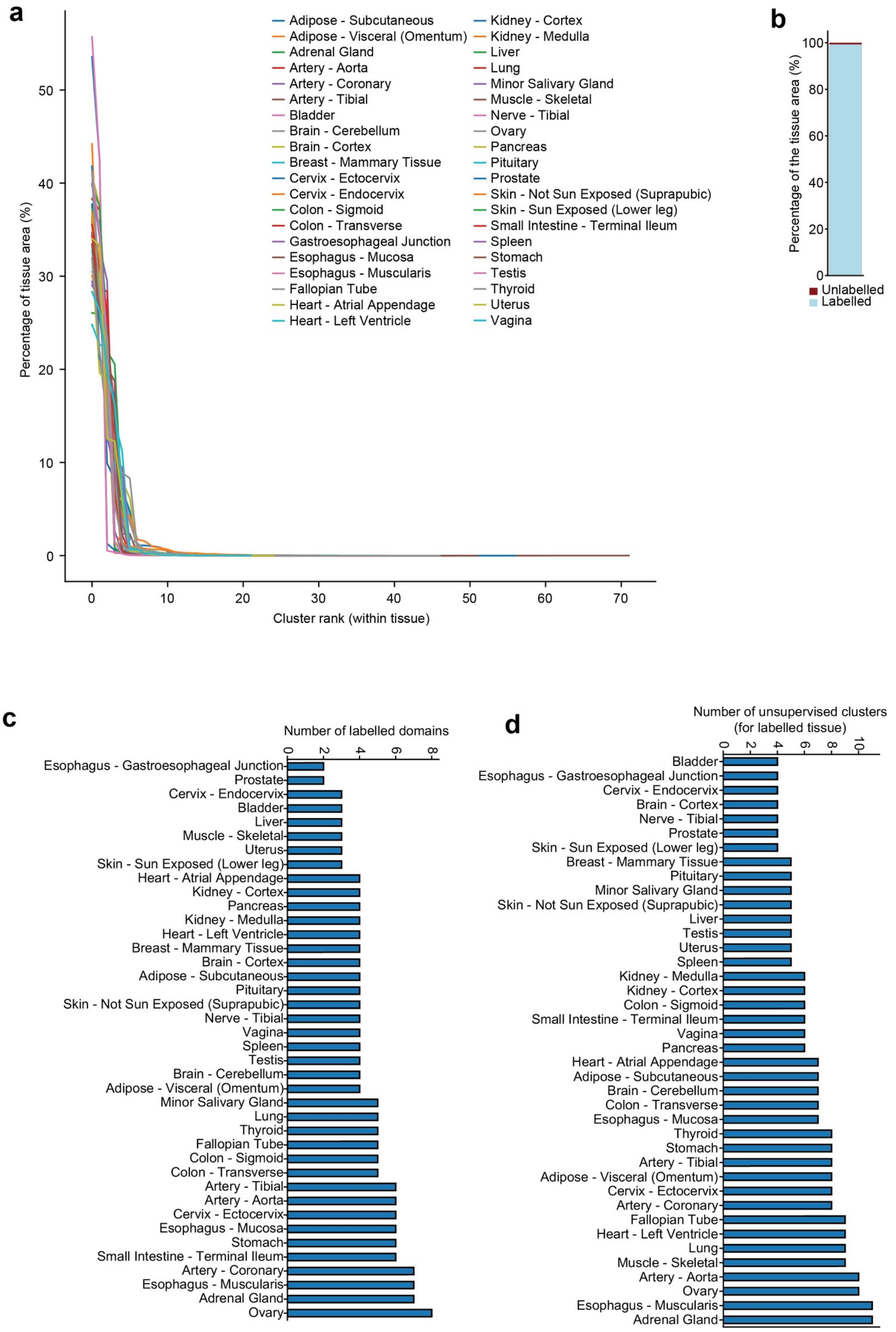


**Supplementary Figure 3. Domain labeling and unsupervised cluster characterization**. **a,** Distribution of tile area per cluster within each tissue. For all tissues, a few clusters dominate the tissue area while the remaining clusters contribute only marginally. **b,** Percentage of labeled vs. unlabeled tissue area across the entire dataset. Over 99% of the analyzed tissue area is associated with an annotated domain label. **c,** Number of unique labeled domains per tissue type. **c,** Number of unsupervised clusters per tissue underlying the labeled domain names.

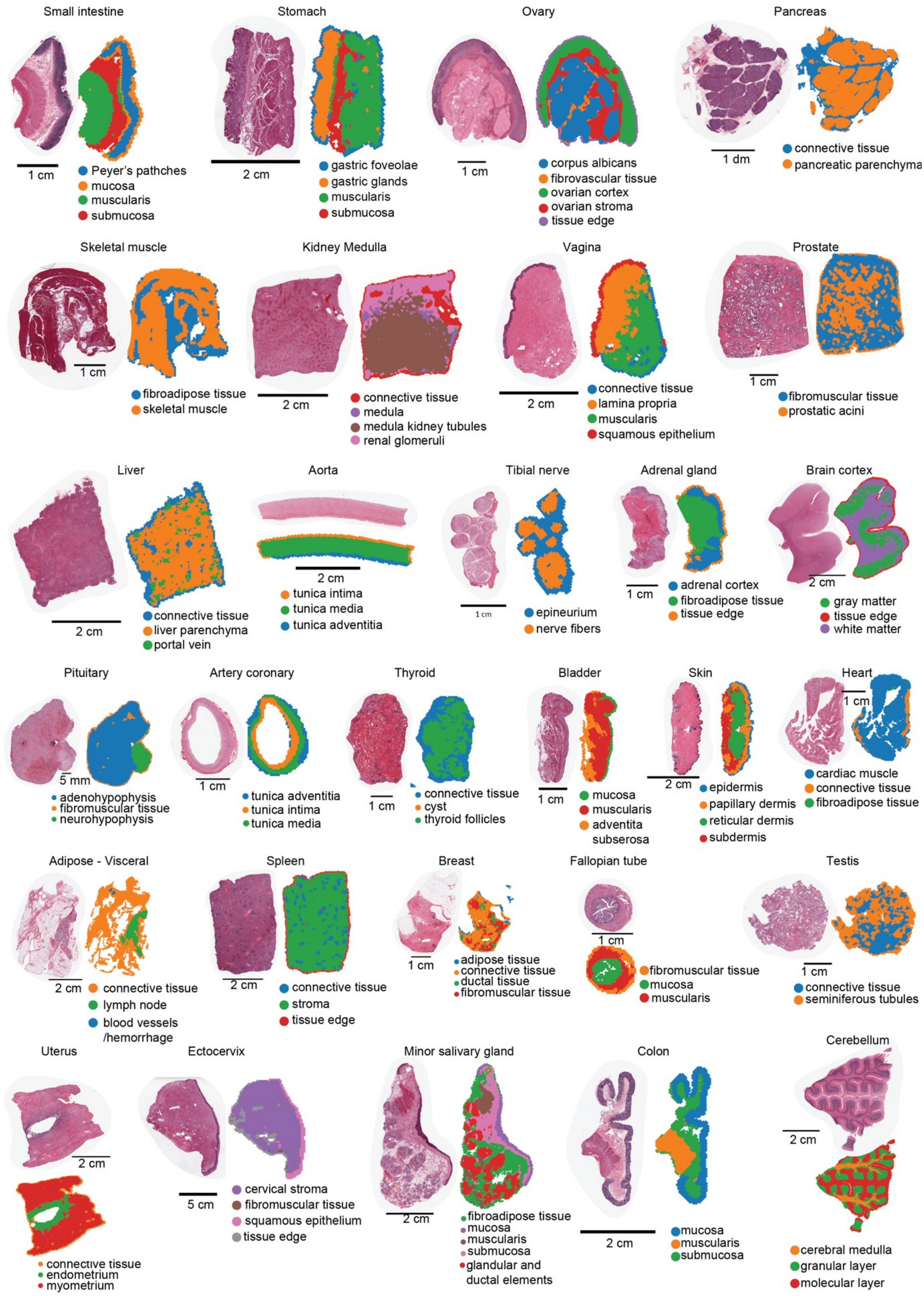


**Supplementary Figure 4. Examples of microanatomical domains in the GTEx dataset.** The colors labeling domains are not matched across tissues.

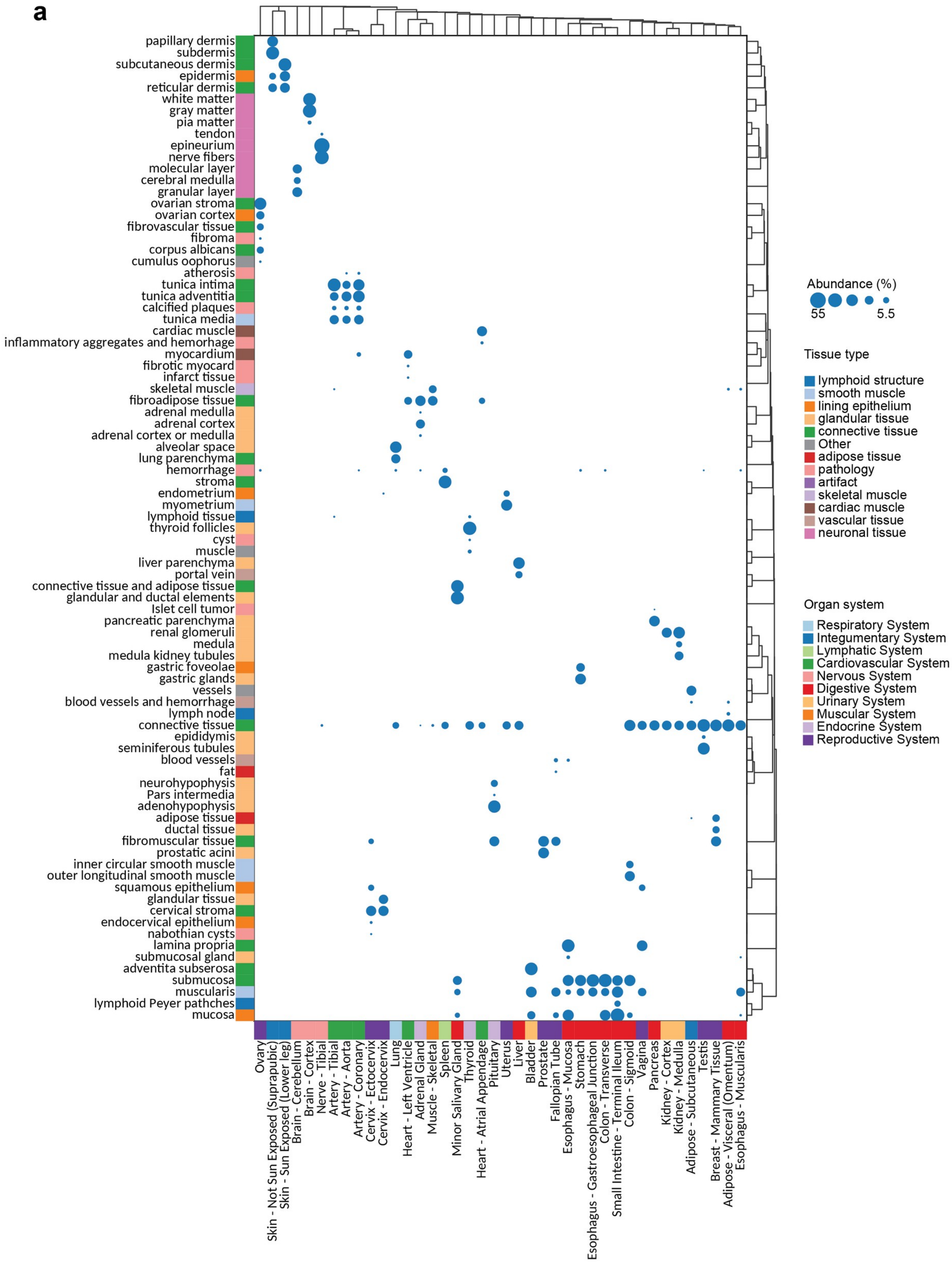


**Supplementary Figure 5. Microanatomical composition of the human body. a,** Abundance of microanatomical domains across tissues.

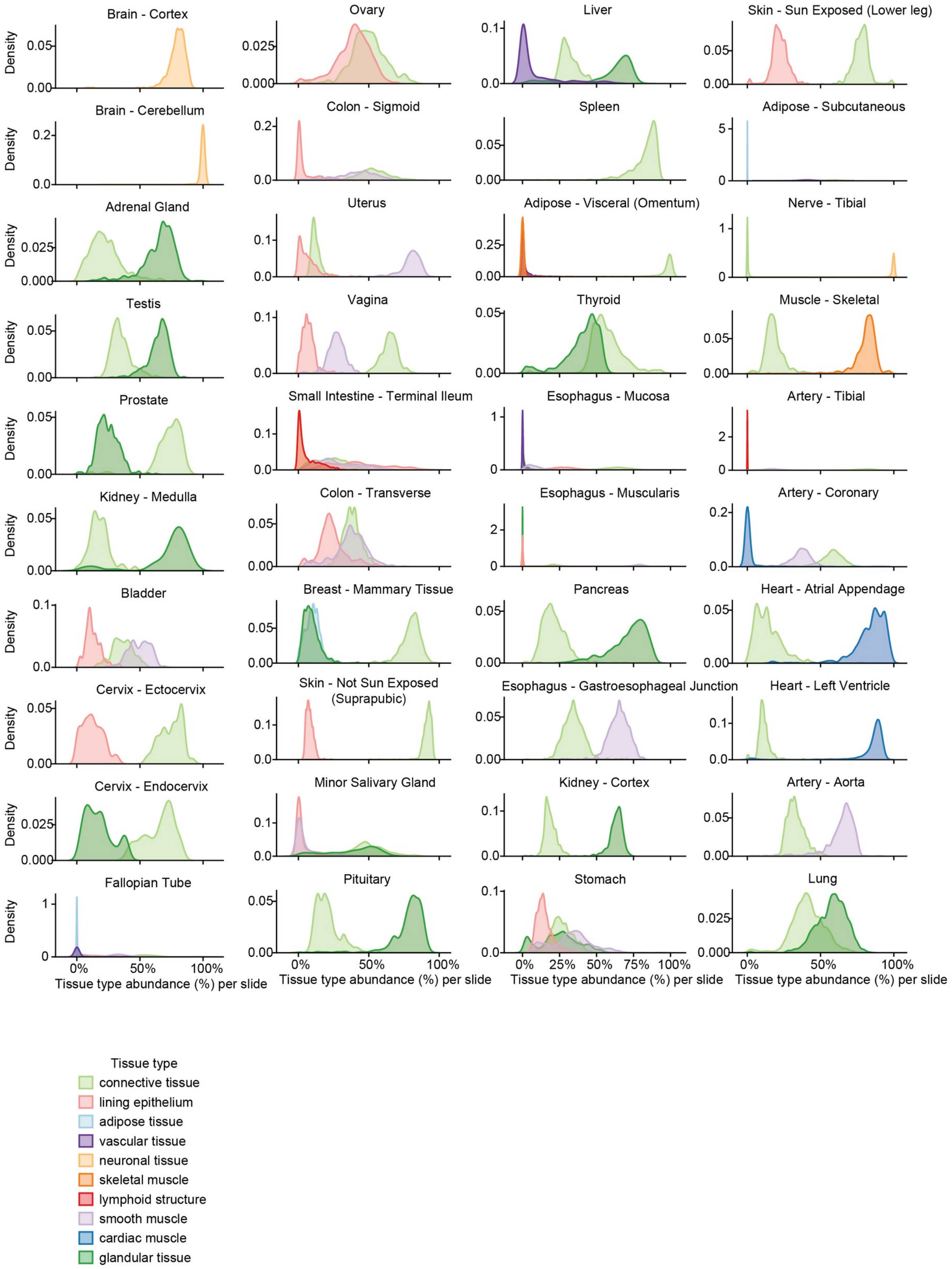


**Supplementary figure 6. Distribution of tissue-type abundances across human tissues.** Density distributions of cluster-level tissue-type abundances per slide, expressed as percentages for all analyzed tissues. Artifacts, pathology and unannotated domains were excluded.

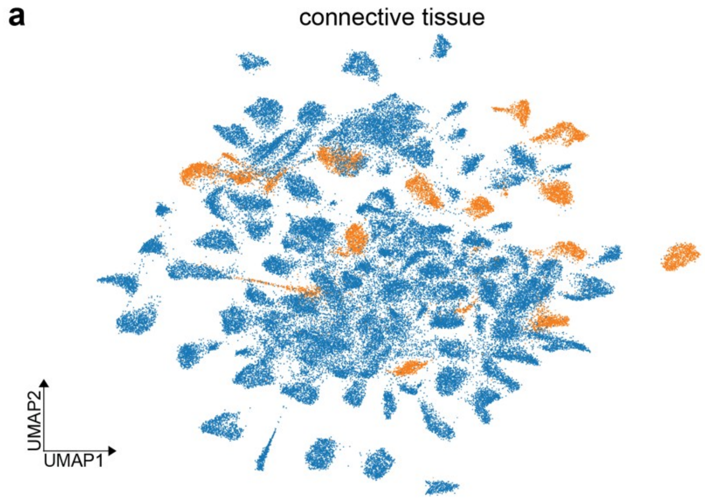


**Supplementary Figure 7. Morphological profile of connective tissue across the dataset.**

**a**

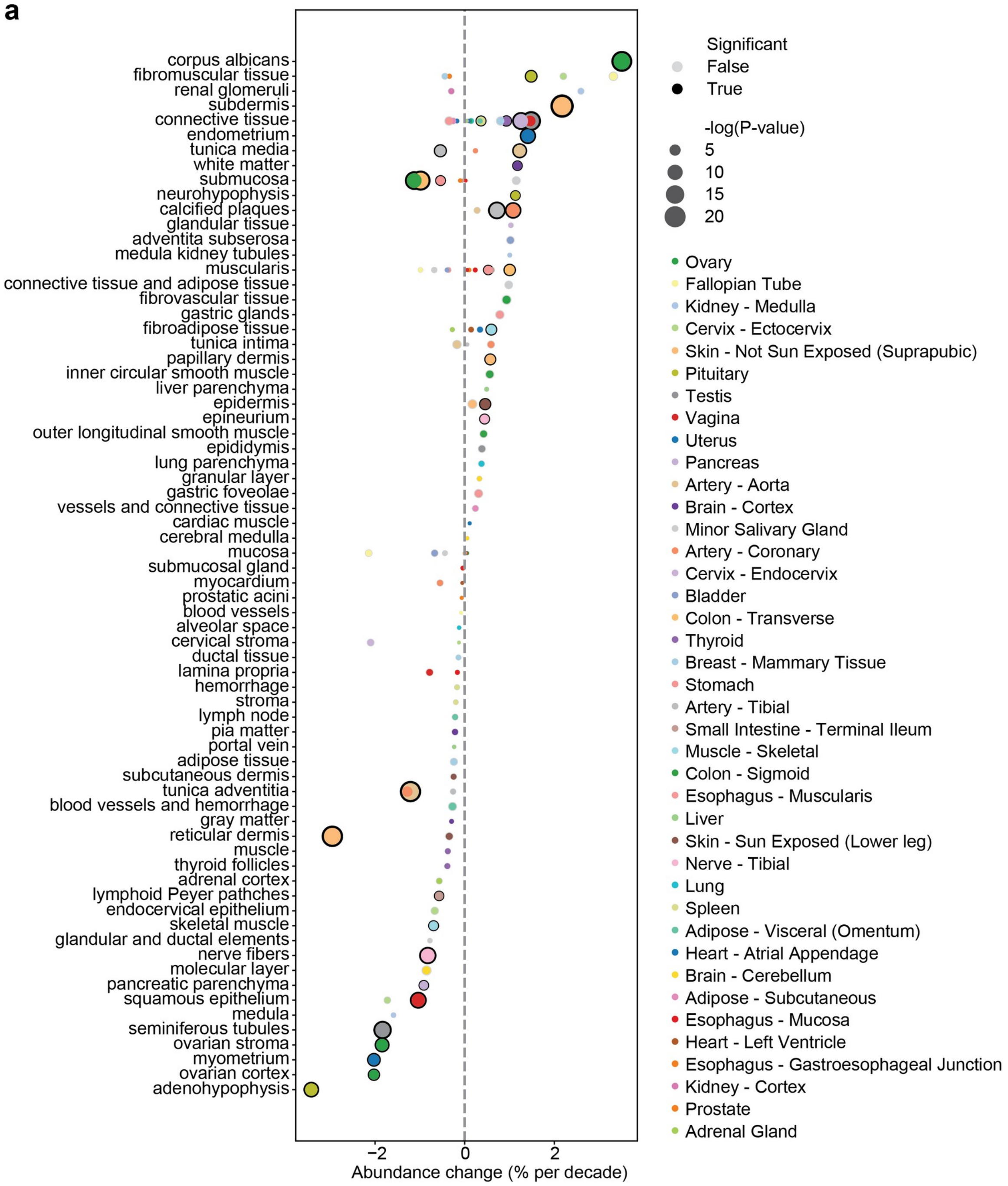


**Supplementary Figure 8. Age-associated changes in microanatomy.** Scatterplot of changes in the abundance of microanatomical domains with age across tissues in the GTEx dataset.

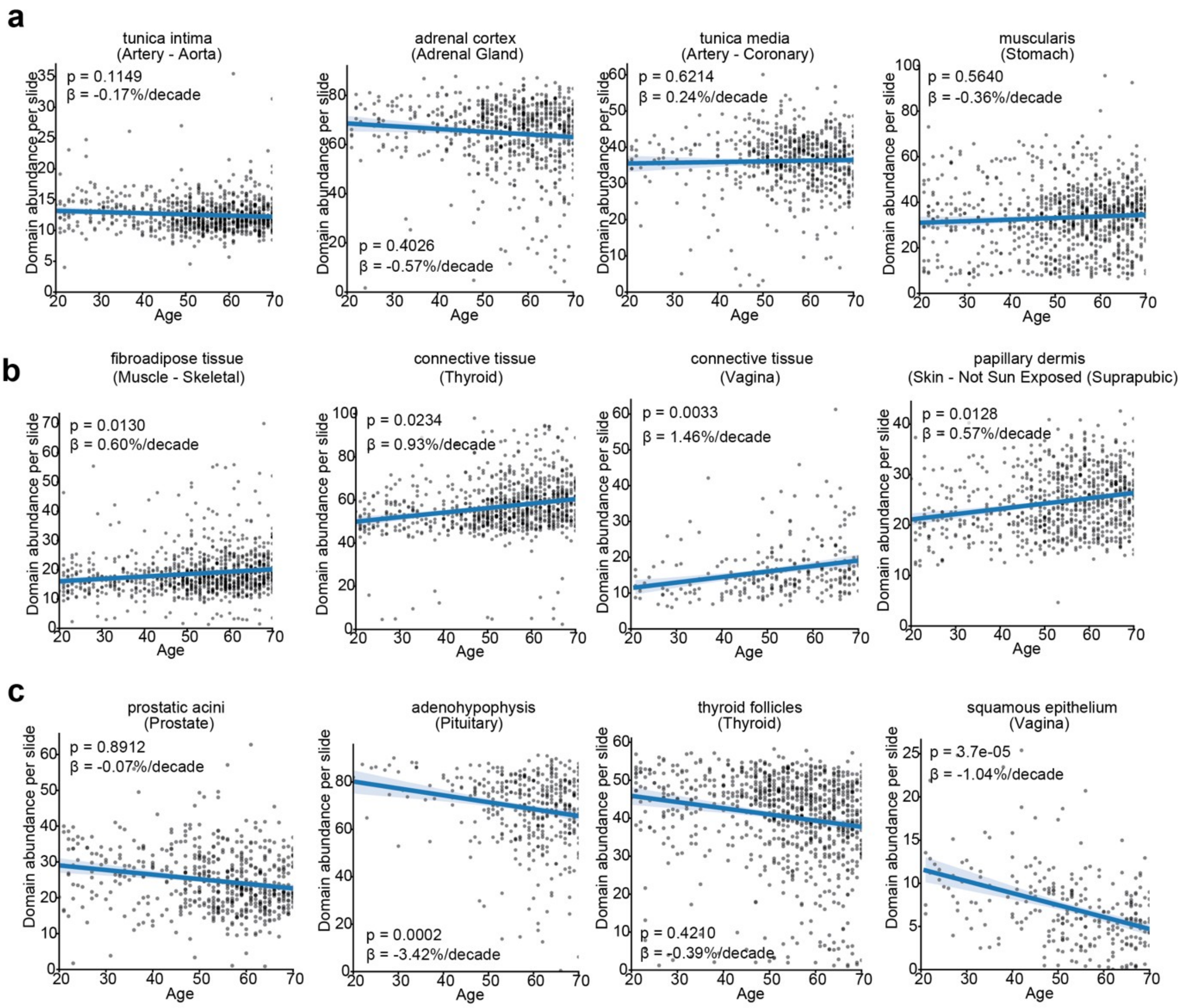


**Supplementary Figure 9. Examples of abundance changes of microanatomical domains with age** for: **a,** domains stable with age; **b,** domains increasing in abundance with age and **c,** domains decreasing in abundance with age. Slopes represent linear regression between age and domain abundance. P-values and coefficient β are derived from linear regression corrected for covariates such as ischemic time, cohort and sex.

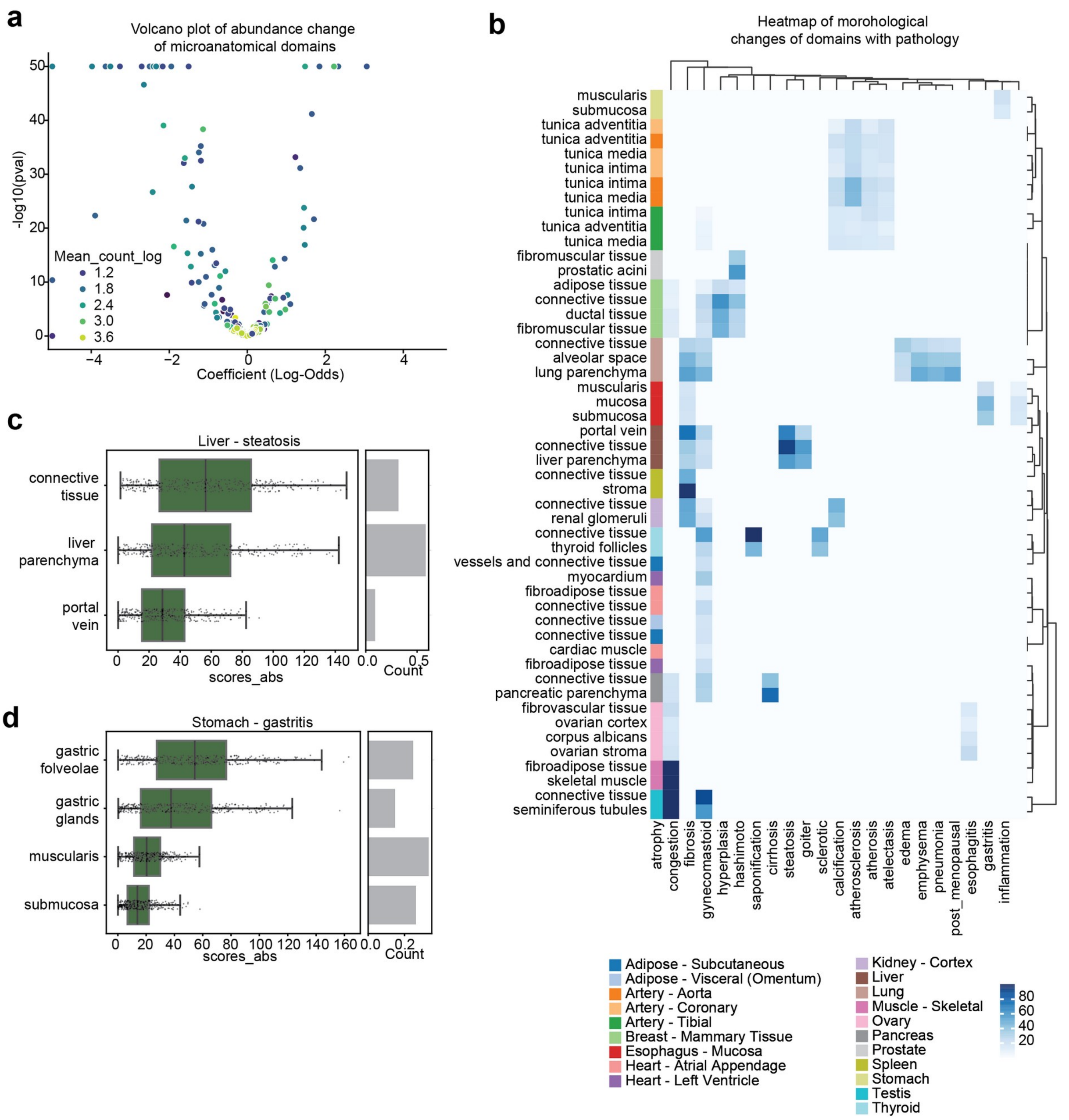


**Supplementary Figure 10. Effects of subclinical pathology on tissue microanatomy. a,** Volcano plot showing changes in the abundance of microanatomical domains associated with subclinical pathology. Each point represents a domain, with effect size on the x-axis (GLM model coefficient) and significance (-log10(p-value)) on the y-axis, colored by the log-transformed mean tile count across samples. **b,** Heatmap displaying the average differential score (a composite metric combining effect size and statistical significance) of morphological features, extracted using the PLIP model, between healthy and pathological samples across tissues. **c,** Example of morphological features showing differential expression across microanatomical domains in steatotic liver. **d,** Example of feature differences across microanatomical domains in gastritis-affected stomach tissue.

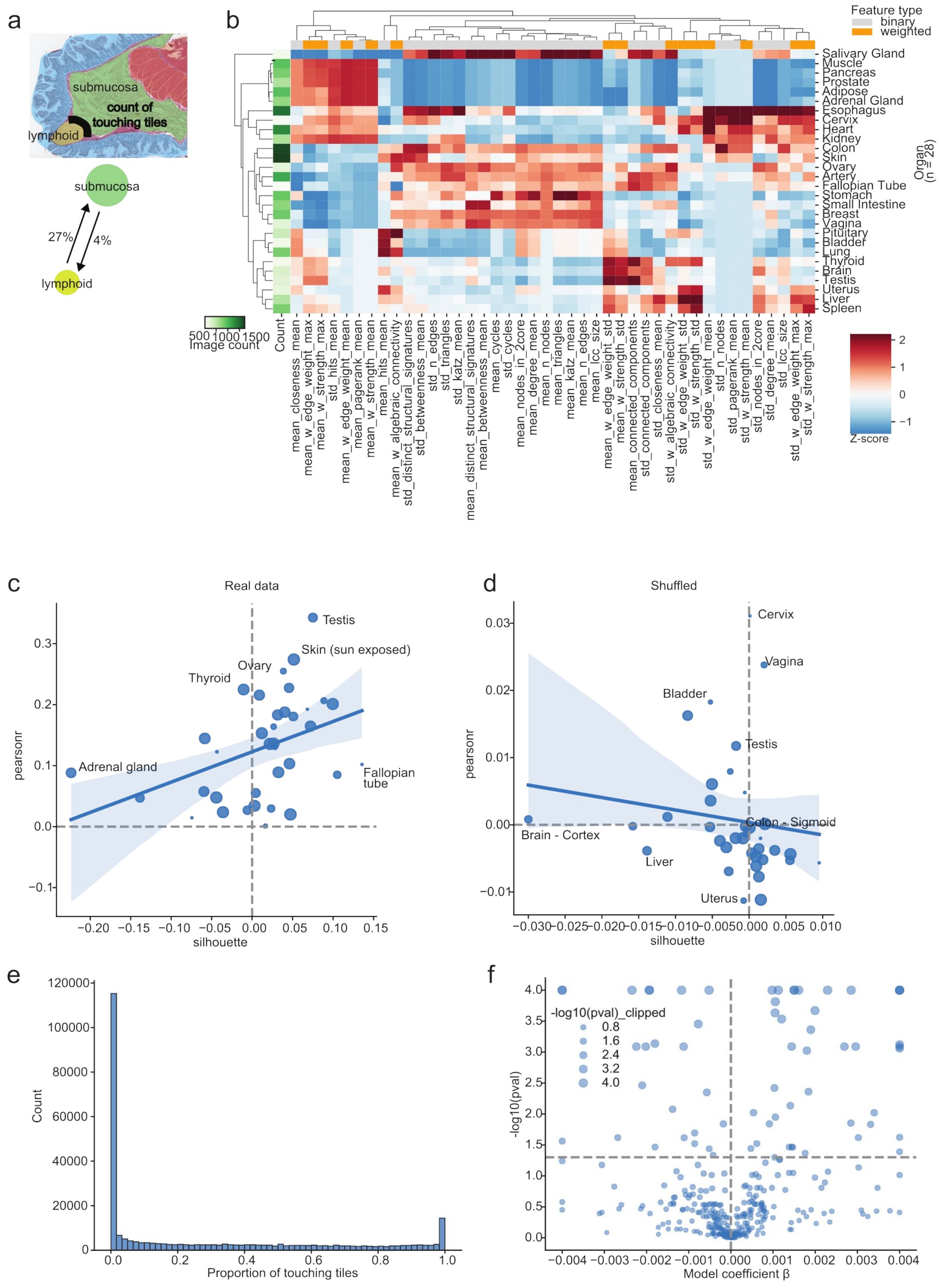


**Supplementary Figure 11. Topological characteristics of tissue microanatomical graphs. a,** Schematic overview of defining domain interactions by proportion of touching border tiles between microanatomical domains. **b,** Heatmap of standardized (z-scored) graph-derived features across tissue types, including measures such as clustering coefficient, betweenness centrality, and domain-level adjacency metrics. Tissues are hierarchically clustered based on these features. **c,  e,** Distribution of the proportion of domain pairs (tiles) in direct spatial contact across all tissues, showing a bimodal pattern suggestive of distinct architectural regimes. **f,** Volcano plot of coefficients (β) and significance levels (–$\log_{10}$ p-value) from a regression model assessing the association between tissue domain interaction features and age. Dot sizes are scaled by clipped –$\log_{10}$(p) values.

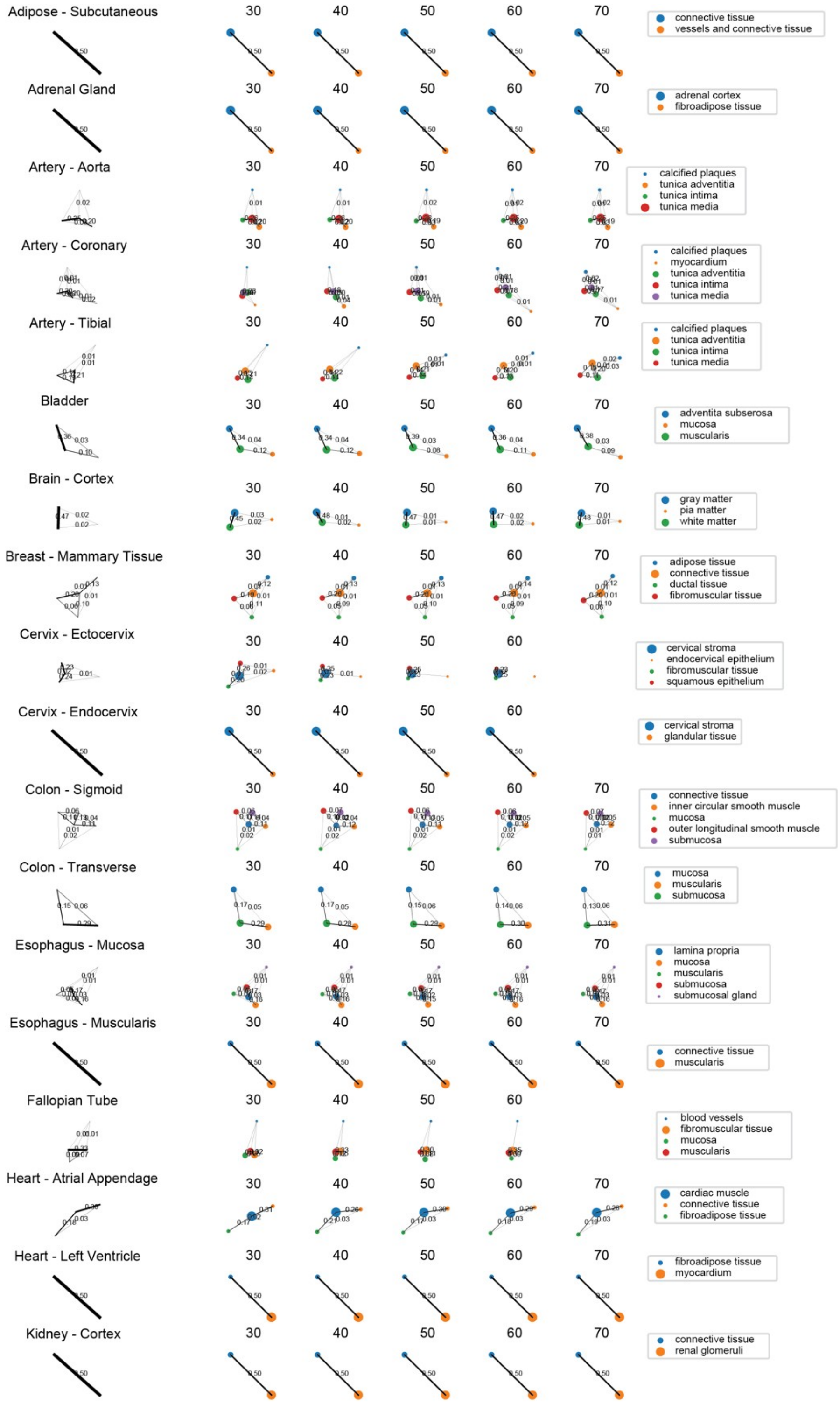


**Supplementary Figure 12. Age-associated changes in domain interaction networks across tissues (Part 1).** Graph representations of domain–domain interaction networks in selected tissues, stratified by age bracket (30, 40, 50, 60, 70 years). Each node represents a microanatomical domain, and edges represent spatial interactions between domains. Edge thickness and opacity are scaled by interaction strength (e.g., frequency or probability). The average (age-aggregated) interaction network for each tissue is shown on the left, with age-stratified graphs in adjacent columns. Domain identities are color-coded (legend, right). Continued on Supplementary Figure 13.

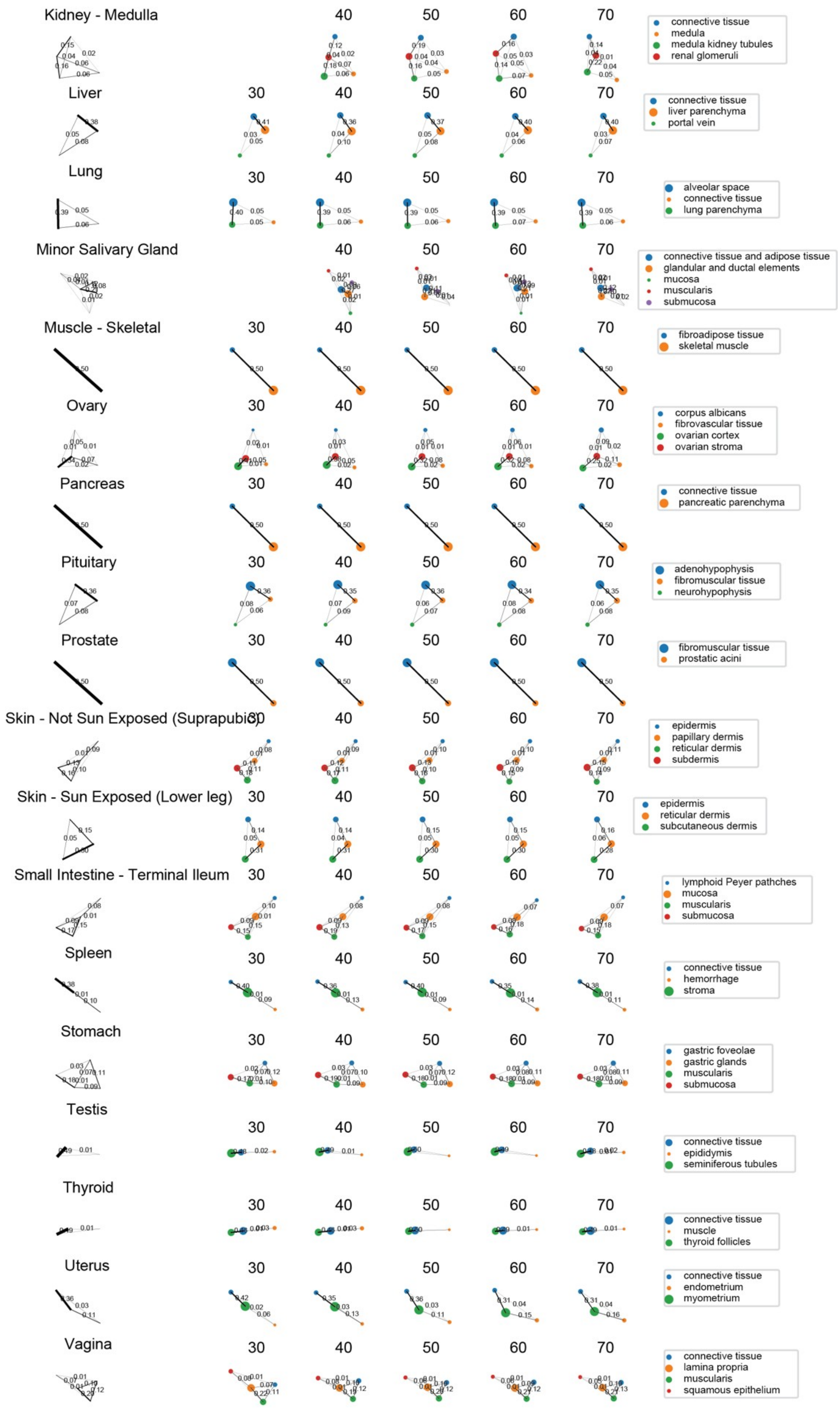


**Supplementary Figure 13. Age-associated changes in domain interaction networks across tissues (Part 2).** Continuation of domain–domain interaction networks for the remaining tissues, stratified by age bracket (30, 40, 50, 60, 70 years). See caption of Supplementary Fig. S12 for description of graphical elements and encoding. These visualizations highlight age-related remodeling of microanatomical architecture in a tissue-specific manner.
